# Genomic conformation reflects epigenetic programming rather than gene expression

**DOI:** 10.64898/2026.11.26.769172

**Authors:** Carol A. Edwards, Frances L. Dearden, Ariella Weinberg-Shukron, Ana Moreno-Barriga, Anjuli K. Freeman, Dionne J. Gray, Anne C. Ferguson-Smith

## Abstract

Genome function is regulated temporally and tissue-specifically through coordinated interplay between epigenetic modifications and genomic elements, however, the extent to which chromatin conformation shapes or is shaped by epigenetic programming and transcriptional activity in vivo, remains largely unresolved. Through analysis of the well-characterised *Dlk1-Dio3* imprinted domain, we have identified tissue and parent-of-origin specific conformations and taken a genetic approach to determine the hierarchical relationships between conformation, epigenetic state and transcription. Parental-origin specific topological domains identified in fetal brain are dependent on the germline-derived imprinting control region. In the absence of specific CTCF binding at a secondary imprinting control element, the parental origin-specific CTCF-mediated conformation can be disrupted without a change to domain-wide imprinting, transcription or epigenetic state. These findings indicate that, in this developmental context, conformation can be decoupled from transcription and that each can be independently governed by a hierarchy of epigenetically controlled regulatory features initiated in the germline.

## Introduction

Genome function is regulated temporally and tissue-specifically through the orchestrated interplay of genomic features and epigenetic states. Recently, attention has turned to the role genome organisation plays in controlling gene expression. At the mega-base level, the genome is organised into topologically associating domains (TADs). TADs are regions of self-interaction that are believed to restrict genes within the same domain to coordinated regional control^1,2^. The boundaries of TADs have been shown to be associated with cohesin and CCCTC-binding factor (CTCF) binding sites: suggesting that they can be formed via loop extrusion of DNA through the cohesin complex until it reaches convergently bound CTCF^3–5^

Although originally thought to be maintained between different tissues, it is now known that some TADs, and especially the sub-TADs and loops (particularly enhancer-promoter interactions) within them, are tissue-specific and orchestrate local genomic contacts throughout development^1,4,6^. The disruption of TAD boundaries has varying effects on transcriptional regulation. For example, deletion of boundary CTCF sites in the *Shh* region alters local conformation but not gene regulation and loss of the whole boundary in the *Sox9-Kcnj2* locus has little effect on expression in the region^7,8^. Conversely, perturbation of two CTCF sites in the alpha-globin locus leads to conformational and transcriptional changes in the region^9^. Furthermore, deletion of four CTCF sites in the *Slc29a3-Unc5b* locus leads to varying, tissue-specific transcriptional responses^10^. Clearly, we are far from fully understanding the relationship between topology and expression. How transcription and topology relate to the epigenetic state of a region also remains largely unknown. A recent study found that increased methylation levels in *Tet* triple knock-out mouse embryonic stem cells (mESCs) resulted in weakening of the TAD structures and loss of long-range loops, but how this relationship functions in vivo is yet to be fully explored^11^.

Genomic imprinting provides a perfect paradigm for studying the role of conformation in genome function as two homologous chromosomal regions, which share the same cellular milieu, express a different repertoire of genes in a parental-origin-specific manner. Imprinted gene expression is established in the germline where DNA methylation is differentially acquired between sperm and oocytes at specific elements called imprinting control regions (ICRs)^12^. ICRs are genomic elements that dictate the monoallelic expression of all imprinted genes within a domain. Using diploid chromatin conformation capture (Dip-C) in the mouse brain, parental-specific conformations have been identified in eight imprinted regions, indicating differential conformation may also be a feature of all imprinted domains^13^. The *Igf2-H19* region has long been known to form parent-of-origin specific conformation^14^. Imprinted gene expression in this domain is controlled by differentially methylated CTCF binding sites in the ICR^15,16^. These CTCF sites are methylated on the paternally inherited chromosome and are thus only able to bind CTCF on the unmethylated maternal chromosome where they function as an insulator preventing *Igf2* from being expressed. In ESCs, the ICR has been shown to form a maternal chromosome-specific sub-TAD boundary that splits the imprinted domain in two^14,17^.

The *Dlk1-Dio3* region is one of the largest imprinted domains in mammals, which has previously been shown to reside within a single TAD in various tissues including ESCs, cortex and liver^1,18,19^. The region contains four protein-coding genes, *Dlk1*, *Rtl1*, *Dio3* and one isoform of *Begain*, that are expressed from the paternally inherited chromosome (Fig. 1A). *Gtl2* (also known as *Meg3*), and its associated non-coding transcripts, are exclusively expressed from the maternally inherited chromosome (Fig. 1A). Imprinting in the domain is controlled by a germline derived, intergenic differentially methylated region (IG-DMR) which acts as the ICR^20^. Maternal deletion of the IG-DMR leads to paternalization of the maternal chromosome and mice die between E16.5 and birth^20^. Recently, we and others have shown that the IG-DMR is a bipartite element consisting of a CpG island (IG-CGI) and a transcription regulatory element (IG-TRE) that is believed to act as an enhancer for *Gtl2* on the unmethylated maternal chromosome^21–24^. Interestingly, deletion of the IG-CGI only is lethal on paternal inheritance, causing the maternalization of the paternal chromosome, but has no effect when deleted from the maternal chromosome^24^.

**Fig. 1.**
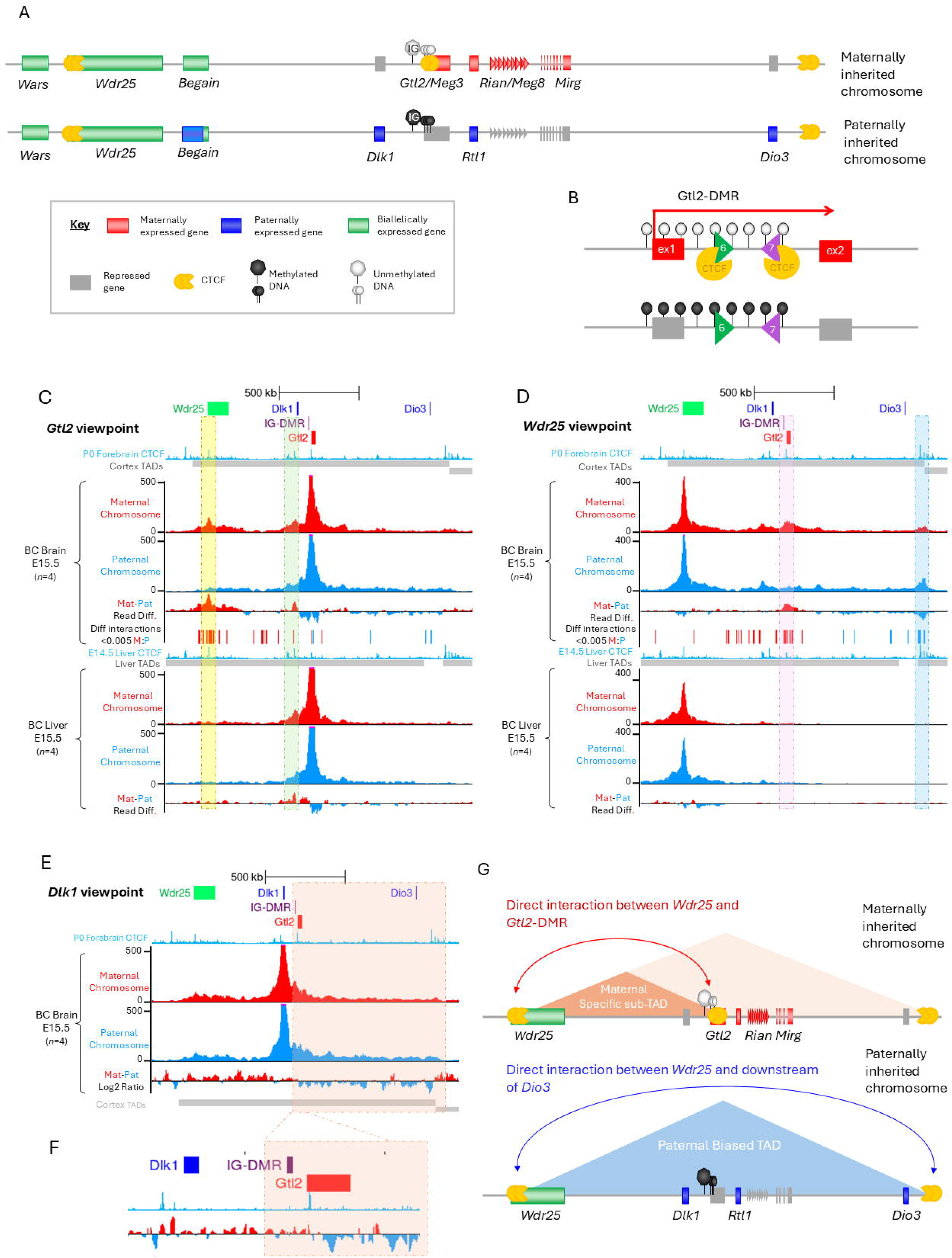
The Dlk1-Dio3 region forms parent-of-origin and tissue-specific conformations in vivo. (**A**) Schematic of the *Dlk1-Dio3* domain. The lncRNAs *Gtl2/Meg3, Rian/Meg8, Mirg* and an antisense transcript of *Rtl1* are exclusively expressed from the maternally inherited allele. Three protein coding genes are expressed from the paternally inherited chromosome: *Dlk1*, *Rtl1* and *Dio3*. An alternative transcript of *Begain* is also predominantly paternally expressed. The ICR for the region is the IG-DMR, which is methylated on the paternal chromosome (black heptagon) but remains unmethylated on the maternal one (white heptagon). (**B**) the Gtl2-DMR contains two CTCF binding sites: forward facing CTCF6 (green arrowhead) and backward facing CTCF7 (purple arrowhead). The region is methylated on paternally inherited chromosome, so CTCF only binds on maternally inherited one. (**C-D**) 4Cseq data in E15.5 fetal brain (upper panel) and fetal liver (lower panel) with the *Gtl2* promoter (**C**) or *Wdr25* intron 2 (**D**) as the viewpoint fragment in C57BL/6 x CAST/EiJ (BC) hybrid mice. (**E**) 4Cseq data with Dlk1 promoter as the viewpoint in E15.5 fetal brain in BC hybrids. (**F**) Zoom in on “switch-point” at Gtl2-DMR. For all 4Cseq data the mean average reads per fragment (normalised to total assigned reads) are shown. Contacts on the maternally inherited chromosome are shown in red and contacts on the paternally inherited chromosome are shown in blue. For Gtl2 (**C**) and Wdr25 (**D**) viewpoints the read difference between the maternal (red) and paternal (blue) chromosome are shown below. For the Dlk1 viewpoint data (**E,F**) the log_2_ ratio of maternal (red) to paternal (blue) contacts are plotted below. Forebrain and E14.5 liver CTCF signal (turquoise) are taken from ENCODE track hub ^30^. All data are viewed on the UCSC genome browser^31^. (**G**) Proposed model of parent-of-origin specific conformations in fetal brain. On the maternally inherited chromosome CTCF can bind to the unmethylated *Gtl2*-DMR. This facilitates a direct interaction to form between the *Gtl2* promoter and intron 2 of *Wdr25*. The CTCF bound *Gtl2*-DMR acts as an insulator, restricting contacts upstream and forming a maternal chromosome specific sub-TAD. On the paternally inherited chromosome CTCF cannot bind to the methylated Gtl2-DMR. This prevents contacts forming between *Wdr25* and *Gtl2*. Instead, *Wdr25* interacts 117 kb downstream of *Dio3* forming a large paternal specific TAD which encompasses the entire imprinted domain.

The domain contains a secondary somatic DMR, the *Gtl2*-DMR (Fig. 1B)^25^, which covers the promoter, first exon and intron 1 of *Gtl2,* is dependent on the IG-DMR, and is established by embryonic day 6.5^26^. Mouse models deleting the *Gtl2*-DMR on the maternally inherited chromosome recapitulate the full ICR deletion with the downregulation of maternally expressed genes, upregulation of paternally expressed genes and embryos dying *in utero*^27,28^. These data emphasise the hierarchical relationship between epigenetically controlled regulatory features at the locus.*Gtl2* intron1 contains two CTCF binding sites (CTCF6 and CTCF7) that are only able to bind CTCF on the unmethylated maternal copies (Fig. 1B). The CTCF6 motif is forward-facing and likely to form interactions downstream, whereas the CTCF7 motif is reverse facing. Previous data using stem cells in culture, indicate that deletion of CTCF6 has no effect on expression or conformation in vitro whereas deletion of CTCF7 leads to a reactivation of maternal *Dlk1* in neural stem cells and loss of conformation^17^. These CTCF binding sites were also shown to coincide with a maternal chromosome-specific sub-TAD boundary in ESCs that divides the previously identified TAD on the maternally, but not the paternally, inherited chromosome^17^. Furthermore, hypo-methylation of the paternal CTCF binding sites in vitro leads the region adopting a maternal-like conformation but it is not clear whether this is a cause or a consequence of transcriptional changes^29^. These findings suggest that, at least in stem cells, CTCF binding at this locus can influence conformation and parental-origin specific transcription and emphasise the value in deciphering the hierarchical relationships between conformation, epigenetic state and transcriptional outcome in vivo.

Here, we use the *Dlk1-Dio3* region as a model to study parental-origin and tissue-specific topology in vivo. By generating a series of mouse deletion models, we have been able to investigate the exquisite interplay between genomic elements, epigenetic marks, expression, and conformation. We find that the region forms both parent-of-origin and tissue-specific conformations in vivo. Furthermore, whilst parent-of-origin expression and conformation are both dictated by epigenetic states conferred in the germline, expression and conformation are decoupled from each other. These findings reveal that the germline epigenetic state independently confers both conformation and transcription and that conformation is neither shaped by nor is shaping transcriptional outcome at this imprinted domain.

## Results

### The *Dlk1-Dio3* region forms parent-of-origin and tissue-specific conformations in vivo

To investigate whether the *Dlk1-Dio3* domain forms parent-of-origin specific conformation during development in vivo, we used 4Cseq (circular chromatin conformation capture with sequencing^32,33^) with the viewpoint fragment in the Gtl2-DMR region. For these experiments, reciprocal hybrids of two mouse strains (C57BL/6J and *Mus musculus castaneus* - CAST/EiJ) were utilised to allow single nucleotide polymorphisms (SNPs) to identify interactions with the maternal and paternal viewpoint.E15.5 brain and liver were chosen as they show reciprocal tissue-specific expression patterns for *Dlk1* and *Gtl2*. In fetal brain *Gtl2* is highly expressed with low levels of *Dlk1* expression, whilst in liver, *Dlk1* expression is high with modest levels of *Gtl2* expression (Supplementary Fig. S1A). In both fetal brain and liver, a maternally inherited chromosome specific interaction is observed between *Gtl2* and *Dlk1* (Fig. 1C - green box). Interestingly, a maternal chromosome-specific interaction is observed between *Gtl2* and the second intron of the biallelically expressed gene *Wdr25* that is absent on the paternally inherited chromosome. This interaction is present in fetal brain but is absent in liver indicating this long-range contact is tissue-specific (Fig. 1C, Supplementary Fig. S1B – yellow boxes). Differential interactions between the maternal and paternal chromosome were called using 4C-ker^34^ and show an enrichment of maternal-chromosome specific interactions in *Wdr25*. This brain-specific interaction corresponds with two forward-facing CTCF binding sites in *Wdr25* intron 2, which are conserved in eutherian mammals, as well as putative enhancer elements as denoted by H3K27ac enrichment (Supplementary Fig. 1F).

Using *Wdr25* intron 2 as the viewpoint to confirm this, the interaction between *Wdr25* and *Gtl2* is again observed on the maternally inherited chromosome in the fetal brain from reciprocal crosses, peaking at the *Gtl2* CTCF binding sites (Fig. 1D, Supplementary Fig. S1C and G – pink boxes). This interaction is absent on the paternally inherited chromosome; instead, a much stronger contact is present between *Wdr25* and 117kb downstream of *Dio3*, which corresponds to the distal edge of the previously defined cortical TAD (Fig. 1D, Supplementary Fig. S1C turquoise boxes)^1^. 4Cseq experiments using the *Dlk1* promoter as the viewpoint show similar interaction patterns on both chromosomes, with most contacts restricted to within the previously defined TAD (Fig. 1E, Supplementary Fig. S1D). However, plotting the log_2_ ratio of the interactions demonstrates that a switch from maternally biased to paternally biased contacts occurs at the Gtl2-DMR, indicating the location of an allele-specific sub-TAD boundary, where CTCF binding to the maternal-chromosome restricts contacts to upstream of the binding sites (Fig. 1F, Supplementary Fig. S1E - orange boxes)

Together, these data illustrate that the *Dlk1-Dio3* domain forms parent-of-origin and tissue-specific conformations in vivo. In the fetal brain, the paternal chromosome forms a large TAD between *Wdr25* and *Dio3*, covering the entire imprinted domain and coinciding with a previously defined TAD^1^, whereas the maternal chromosome forms a smaller sub-TAD between the Gtl2-DMR and *Wdr25* (Fig. 1G). This interaction is probably anchored by the two, forward-facing, CTCF binding sites in *Wdr25* and CTCF7 in the Gtl2-DMR, which is reverse-facing and only binds CTCF on the unmethylated maternal chromosome. In contrast, most liver contacts are restricted to much smaller regions, within 250kb of the viewpoints, forming discrete contact regions within a previously determined 2.2Mb TAD that spans from ∼40kb upstream of *Bcl11b* to ∼40kb upstream of *Dio3* (Fig. 1C,D)^18^

Next, we explored whether this parent-of-origin specific conformation is dictated by epigenetics and/or gene expression using a systematic series of genetic mutations that can distinguish these three parameters.

### Parent-of-origin-specific conformation in fetal brain is lost upon deletions of and within the imprinting control region

We first investigated conformation when the epigenotype of the maternal chromosome is switched to a paternal-like state by deleting the IG-DMR. To discriminate between the two chromosomes, IG-DMR deletion mice^20^ were crossed to CAST/EiJ wildtypes in reciprocal crosses. Using the *Gtl2* promoter as the viewpoint, the maternal chromosome-specific interaction with *Wdr25* is absent in maternal IG-DMR heterozygotes (Fig. 2B – yellow box). Consistent with this, when *Wdr25* is the viewpoint, the interaction between *Wdr25* and *Gtl2* is lost, and the interaction downstream of *Dio3* is stronger at a level comparable to the paternal chromosome (Fig. 2C – pink and turquoise boxes). Finally, using *Dlk1* as the viewpoint and plotting the log_2_ ratio of maternal and paternal contacts, the switch from predominantly maternal contacts upstream of Gtl2-DMR to predominantly paternal contacts downstream of this region seen in wildtypes, is missing in the maternal IG-DMR heterozygotes, indicating loss of the maternal chromosome-specific subTAD boundary at the Gtl2-DMR (Fig. 2D – orange box). Upon paternal transmission of the IG-DMR deletion, the paternal chromosome-specific conformation is unchanged (Supplementary Fig. S2A-C). Thus, when the maternal chromosome undergoes a switch to behave epigenetically and transcriptionally as a paternal chromosome, it also adopts a paternal chromosome-specific fetal brain conformation (Fig. 2A). Maternal deletion of the IG-DMR had no impact on conformation in the fetal liver, suggesting conformation in liver is independent of gene expression (Supplementary Fig. S2E).

**Fig. 2.**
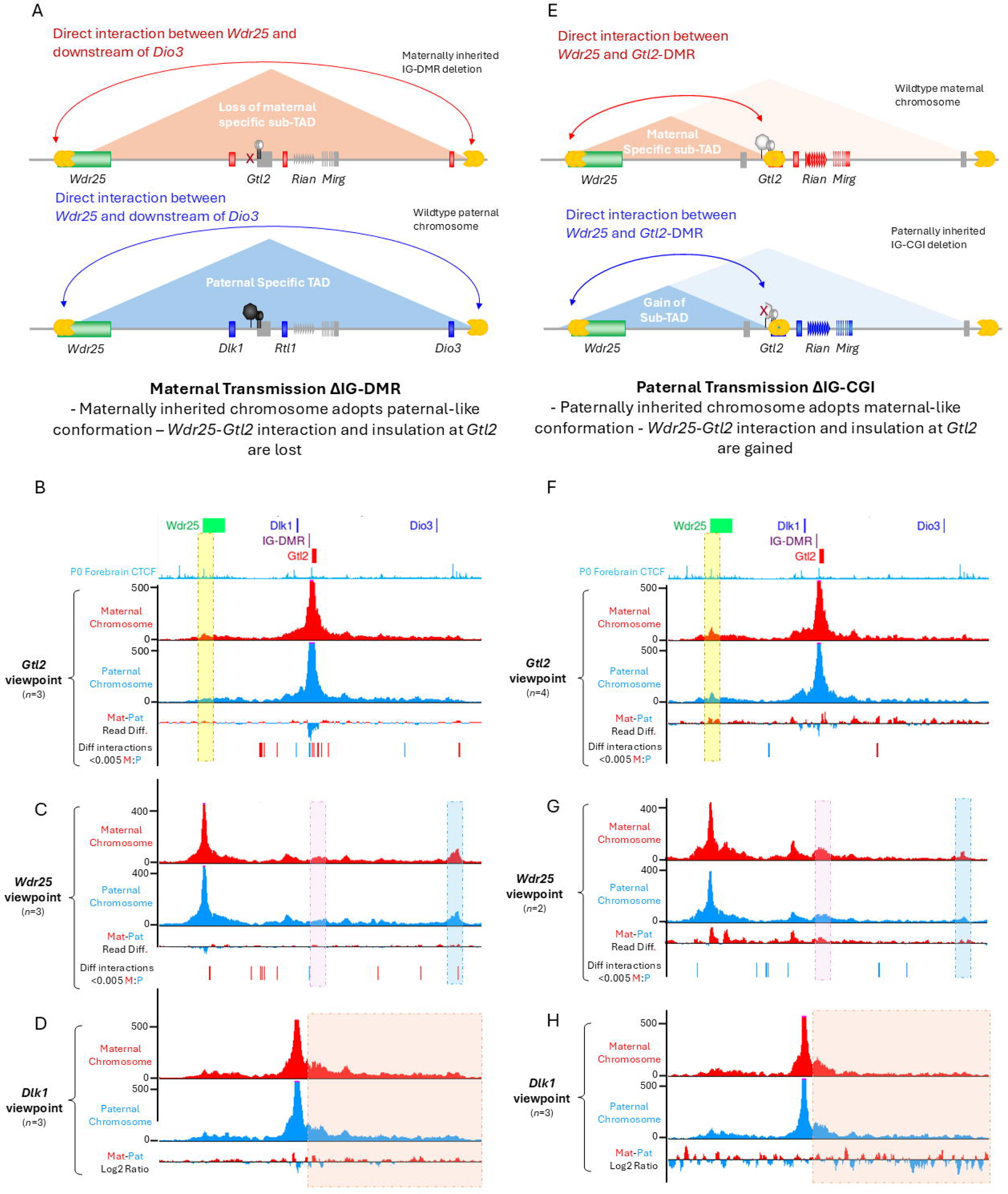
Maternal chromosome specific conformation in fetal brain is lost upon perturbation of the IG-DMR. (**A**) Maternally inherited chromosome adopts paternal-like conformation upon deletion of the IG-DMR. This coincides with a maternal to paternal epigenotype switch, repression of the normally maternally expressed genes and reactivation of paternally expressed genes ^20^. (**B-D**) 4Cseq in fetal brains of IG-DMR deletion maternal heterozygotes (Maternal ΔIG) for the *Gtl2* viewpoint (**B**), *Wdr25* viewpoint (**C**) and *Dlk1* viewpoint (**D**). (**E**) Paternally inherited chromosome adopts maternal-like conformation upon deletion of the IG-CGI. This coincides with a paternal to maternal epigenotype switch, repression of the normally paternally expressed genes and reactivation of maternally expressed genes ^24^. (**F-H**) 4Cseq in fetal brains of IG-CGI deletion paternal heterozygote (Paternal ΔIG-CGI) for the *Gtl2* viewpoint (**F**), *Wdr25* viewpoint (**G**) and *Dlk1* viewpoint (**H**). For 4Cseq data, the mean average reads per fragment (normalised to total assigned reads) are shown. Contacts on the maternal chromosome are shown in red and contacts on the paternal chromosome in blue. For Gtl2 and Wdr25 viewpoints the read difference between the maternal (red) and paternal (blue) chromosome are shown below. Significant differential contacts (P<0.005) between the maternal and paternal chromosome were called by 4C-ker^34^ for the *Gtl2* (**B,E**) and *Wdr25* (**C,G**) data: significant maternal contacts are shown in red and significant paternal contacts are shown in blue. For the *Dlk1* viewpoint data (**D,H**) the log_2_ ratio of maternal (red) to paternal (blue) contacts are plotted below.

Next, the experiment was repeated using IG-CGI mutants that cause the paternal chromosome to gain a maternal chromosomal epigenotype when the deletion is inherited from the father^24^. Upon paternal transmission of IG-CGI deletion, the paternal chromosome gains the interaction between *Wdr25* and *Gtl2* and the interaction between *Wdr25* and downstream of *Dio3* is reduced (Fig. 2F - yellow box and 2G - pink and turquoise boxes). When *Dlk1* is used as the viewpoint, the switch point is again lost, indicating both chromosomes adopt the same conformation (Fig. 2H – orange box). Therefore, when the paternal chromosome is epigenetically switched to behave as a maternal chromosome it also adopts a maternal chromosome-like conformation (Fig. 2E).

Together, these data show that parent-of-origin specific chromatin conformations observed in the fetal brain are defined by differential epigenetic states that are established in the germline. However, these experiments do not reveal whether the conformational changes are a direct result of the epigenetic state, which in turn drives the transcriptional switch or if the changes in gene expression are directing the conformational differences.

### Maternal deletion of CTCF7 binding site alters conformation without impacting transcription

To further decipher the hierarchical relationship between epigenetics, expression and conformation, we next generated in vivo mouse models that perturb CTCF binding in the Gtl2-DMR using CRISPR-Cas9 (Supplementary Fig. S3.1A-C). Mice harbouring an 11bp deletion across CTCF6 (ΔCTCF6 – Fig. 3A), which results in failure to bind CTCF (Supplementary Fig. S3.1C), were viable and fertile. ΔCTCF6 mice show no loss of imprinting and no changes in expression in genes across the entire domain upon either maternal or paternal transmission of the mutation (Fig. 3B-C, Supplementary Fig. S3.2B-C and Supplementary Fig. S3.3A-D). CTCF6 deletion resulted in no change in DNA methylation (Fig.3D). 4Cseq analysis using the *Gtl2* promoter as the viewpoint shows that maternal or paternal loss of the forward facing CTCF6 site have no effect on conformation (Fig. 3E - upper panel, yellow box).

**Fig. 3.**
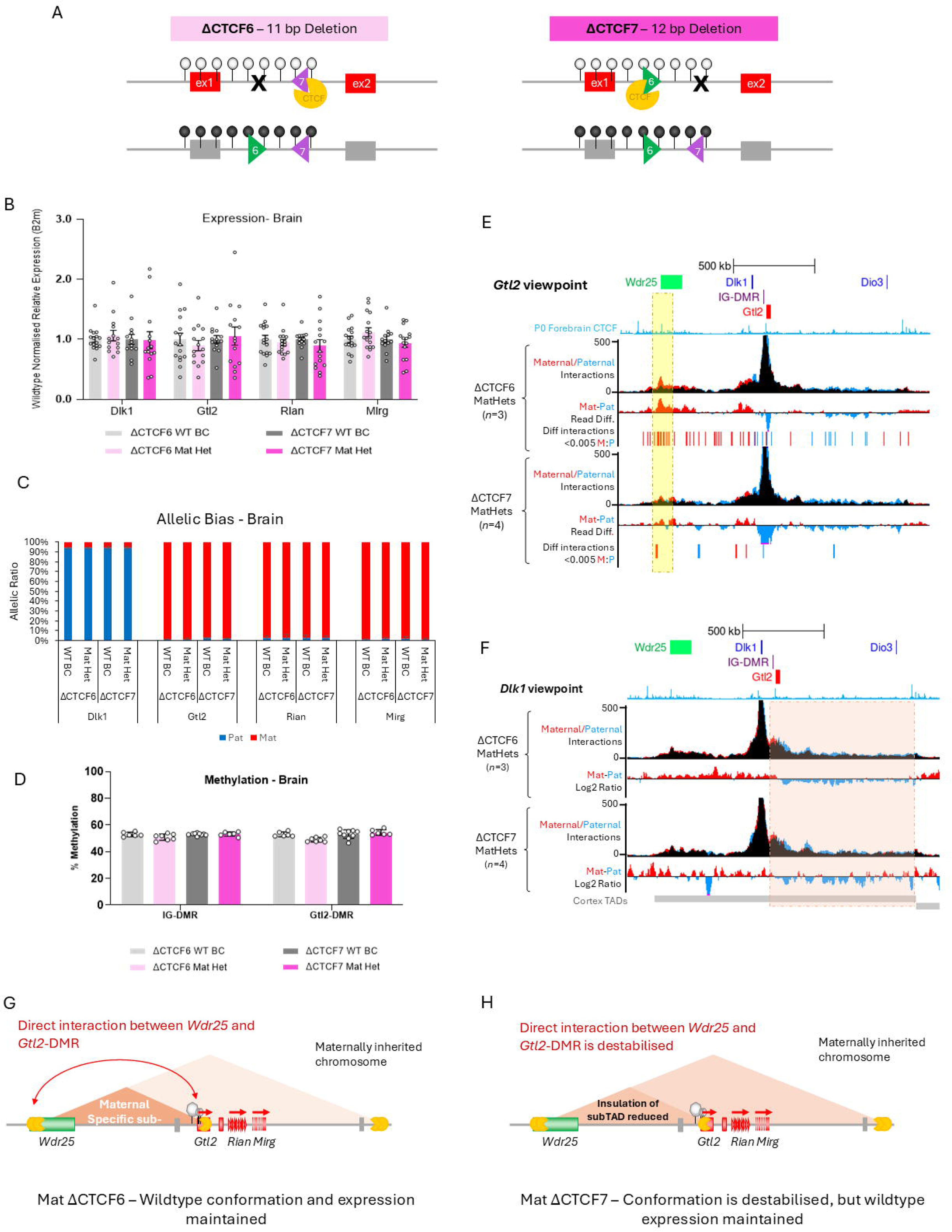
Maternal deletion of CTCF7 binding site alters conformation without impacting transcription. (**A**) Schematic of the Gtl2-DMR in ΔCTCF6 and ΔCTCF7 maternal heterozygotes. (**B**) Expression analysis of ΔCTCF6 and ΔCTCF7 maternal heterozygote E15.5 brains compared with wildtype littermates (WT BCs) as determined by RT-qPCR. Wildtype normalised expression relative to *B2m* is shown. ΔCTCF6 WT BC n = 16, ΔCTCF6 Mat Het n = 14-16, ΔCTCF7 WT BC n = 13, ΔCTCF7 Mat Het n = 14-15, Error bars represent SEM, significance of difference determined using two-tailed non-paired *t* test. (**C**) Ratio of expression from maternal (red) and paternal (blue) allele as determined by pyrosequencing. Error bars represent standard deviation; sample numbers are the same as for expression analysis. (D) Methylation analysis of ΔCTCF6 and ΔCTCF7 maternal heterozygotes in E15.5 brain as determined by pyrosequencing. Regions tested were the IG-DMR, Gtl2 promoter. ΔCTCF6 WT BC n = 6, ΔCTCF6 Mat Het n = 7, ΔCTCF7 WT BC n = 9, ΔCTCF7 Mat Het n = 6. Error bars represent standard deviation, significance of difference determined using two-tailed non-paired *t* test. (**E-F**) 4Cseq in E15.5 fetal brains of ΔCTCF6 and ΔCTCF7 maternal heterozygotes for the *Gtl2* viewpoint (**E**), and *Dlk1* viewpoint (**F**). For 4Cseq data, the mean average reads per fragment (normalised to total assigned reads) are shown. Contacts on the maternal chromosome are shown in red, contacts on the paternal chromosome in blue and overlapping contact are shown in black. For the Gtl2 viewpoint (**E**), the read difference between the maternal (red) and paternal (blue) chromosome are shown below. Significant differential contacts (P<0.005) between the maternal and paternal chromosome were called by 4C-ker ^34^ with significant maternal contacts in red and significant paternal contacts in blue. The *Gtl2-Wdr25* interaction is highlighted in yellow. For the *Dlk1* viewpoint data (**F**) the log_2_ ratio of maternal (red) to paternal (blue) contacts are plotted below. (**G**) In maternal ΔCTCF6 heterozygotes loss of binding at CTCF6 has no impact on gene expression or conformation. (**H**) In maternal ΔCTCF7 heterozygotes loss of binding at CTCF7 destabilises the interaction between *Wdr25* and *Gtl2* and insulation at subTAD boundary is reduced. Loss of CTCF binding at site 7 does not affect gene expression in the domain.

We next investigated a 12bp deletion within CTCF7 (ΔCTCF7 – Fig. 3A) which fails to bind CTCF (Supplementary Fig. S3.1C). Mice are viable and fertile (data not shown). This deletion also had no influence on expression or imprinting across the region upon maternal transmission (Fig. 3B-C, Supplementary Fig. S3.3 A and C). In contrast, on paternal transmission of the CTCF7 deletion, there is a 15% decrease in *Dlk1* expression (p= 0.021) coupled with significant upregulation of *Gtl2*, *Rian* and *Mirg* expression on the paternal chromosome from approximately 2% in wildtypes to 12%, 15% and 17% of total expression, respectively, in paternal heterozygotes (*Gtl2* p=1.21x10^-5^, *Rian* p=3.49x10^-5^ and *Mirg* p=1.34x10^-5^ Supplementary Fig. S3.2A-C). However, although the change to allelic ratio is significant, the total expression levels show a non-significant trend to increased expression of *Gtl2*, *Rian* and *Mirg*, suggesting it is not biologically relevant.

However, despite no change in expression on the maternal chromosome, deletion of the reverse-facing CTCF7 site leads to a destabilisation of the *Gtl2*/*Wdr25* interaction on the maternally inherited chromosome, as demonstrated by the similar peak height and loss of significantly different contacts between the maternal and paternal chromosomes (Fig. 3E – lower panel, yellow box). This indicates that this interaction does not represent a vital enhancer-promoter interaction. In addition, when the *Dlk1* promoter is used as the viewpoint, the tight insulation of the *Gtl2*/*Wdr25* sub-TAD is reduced in maternal ΔCTCF7 brains, with more maternally biased contacts now observed downstream of the Gtl2-DMR (Fig. 3F – lower panel, orange box). Maternal ΔCTCF6 brains maintain this sub-TAD boundary (Fig. 3F – upper panel). No conformational differences were observed on paternal inheritance of the CTCF7 deletion (Supplementary Fig. 3.2D and E).

Therefore, although both CTCF binding site mutations have no influence upon gene expression in the region, loss of CTCF binding at CTCF7 on the maternal chromosome does cause a change in the topology of the domain. This suggests that the tissue-specific contact between *Wdr25* and *Gtl2* is not the result of a promoter/enhancer interaction and that this interaction is not driving the high levels of *Gtl2* expression seen in the brain compared with liver. Together this indicates that conformation is not dictating the expression of this region in the fetal brain.

### *Gtl2* expression is uncoupled from parent-of-origin-specific conformation and *Dlk1* regulation

Next, a mouse line that perturbed CTCF binding at both Gtl2-DMR binding sites was investigated. Δ10Ins6, has a 10bp deletion in CTCF6 and a 6bp insertion in CTCF7. CTCF binding at both sites is disrupted (Supplementary Fig. S3.1C). Mice are viable and fertile (data not shown) but, in contrast to the single site deletion models, this mutation leads to a significant reduction in transcription of the maternally expressed genes on maternal transmission (Fig. 4B, Supplementary Fig. S3.3A). *Gtl2* expression is reduced to 60% of the wildtype level following maternal inheritance of Δ10Ins6 (p= 0.0013). Both *Rian* and *Mirg* expression levels are reduced to 83% of wildtype littermates (p=0.0248 and 0.0183 respectively, Fig. 4B). This downregulation of the maternally expressed genes occurs on the maternal chromosome with no loss of imprinting (Fig. 4C, Supplementary Fig. S3.3C). Furthermore, DNA methylation levels across the IG-DMR, Gtl2-DMR and CTCF binding sites are unchanged, indicating the downregulation is not due to changes at the key epigenetic regulatory elements.(Fig. 4D). *Dlk1* expression was unaffected despite the downregulation of *Gtl2,* suggesting that this level of reduced dosage of the *Gtl2* lncRNA is not directly influencing *Dlk1* levels in the fetal brain. No change in transcription was seen in fetal liver, demonstrating that both CTCF binding sites are required for correct gene expression in a tissue-specific context (Fig. 4E and F). Loss of the two CTCF binding sites on the normally methylated paternal chromosome had no impact on gene expression in the domain (Supplementary Fig. S4B and C, Supplementary Fig. S3.3B and D). Altogether, expression data indicate that CTCF6 and 7 have a combined role in the tissue-specific regulation of maternally expressed genes in the *Dlk1-Dio3* domain.

**Fig. 4.**
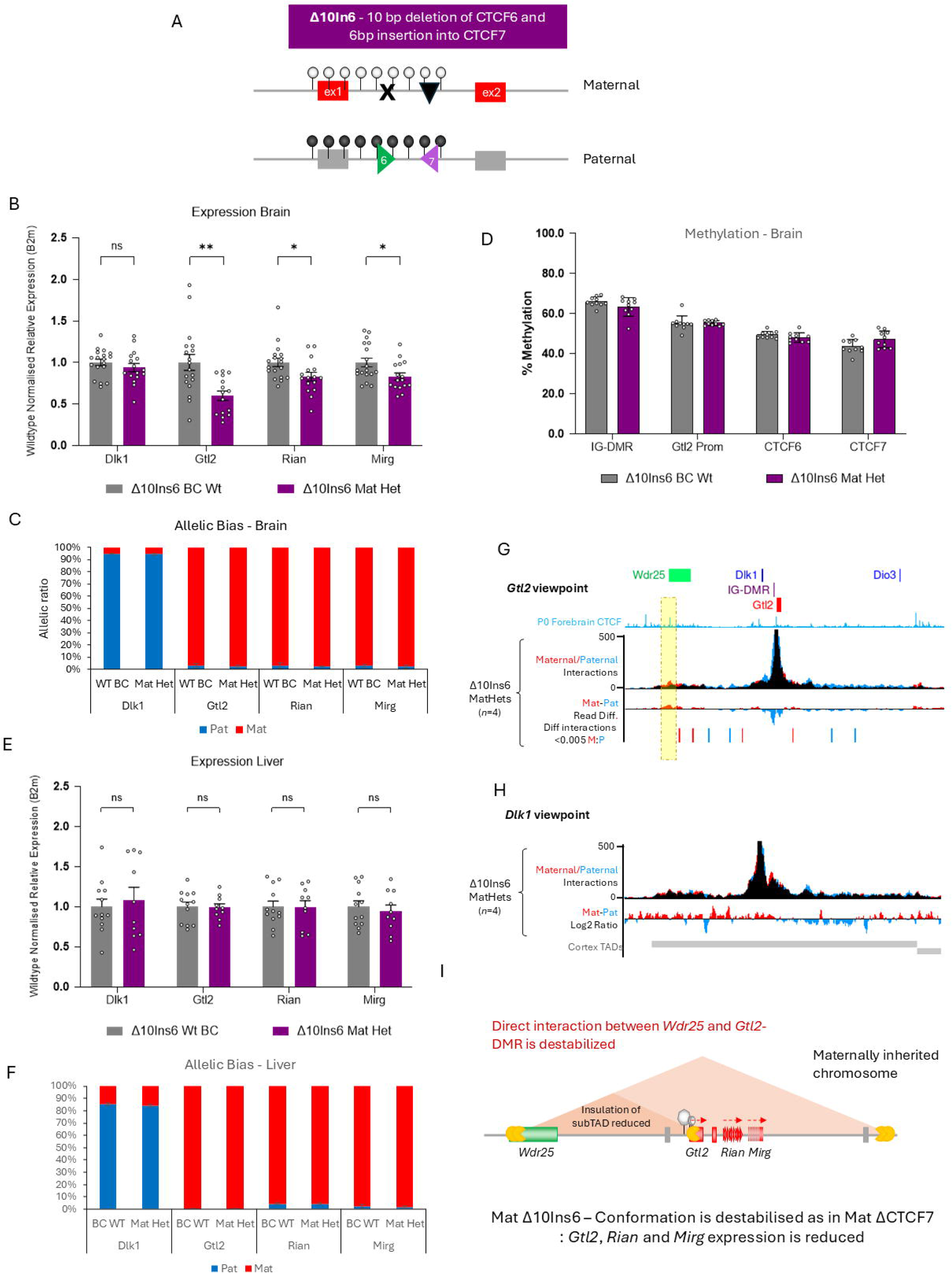
Gtl2 expression is uncoupled from parent-of-origin-specific conformation and Dlk1 regulation. (**A**) Schematic of the Gtl2-DMR in Δ10Ins6 maternal heterozygotes. (**B,D**) Expression analysis in E15.5 brain (B) and liver (D) of Δ10Ins6 maternal heterozygotes compared with wildtype littermates as determined by RT-qPCR. Wildtype normalised expression relative to *B2m* is shown. Brain wildtype n=18, heterozygotes n=15-16 Liver wildtype n=12 heterozygotes n=10. Error bars represent SEM, significance of difference determined using two-tailed non-paired *t* test. (**C,E**) Ratio of expression from maternal (red) and paternal (blue) allele as determined by pyrosequencing in E15.5 fetal brain (**C**) and liver (**E**). Error bars represent standard deviation; sample numbers are the same as for expression analysis. (**F**) Methylation analysis of Δ10Ins6 maternal heterozygotes in E15.5 brain as determined by pyrosequencing. Regions tested were the IG-DMR, Gtl2 promoter region, CTCF6 and CTCF7. Wildtype n=10, heterozygotes n=10. Error bars represent standard deviation, significance of difference determined using two-tailed non-paired *t* test. (**G-H**) 4Cseq in E15.5 fetal brains of Δ10Ins6 maternal heterozygotes for the *Gtl2* viewpoint (**G**), and *Dlk1* viewpoint (**H**). For 4Cseq data, the mean average reads per fragment (normalised to total assigned reads) are shown. Contacts on the maternal chromosome are shown in red, contacts on the paternal chromosome in blue and overlapping contacts are shown in black. For the *Gtl2* viewpoint (**G**), the read difference between the maternal (red) and paternal (blue) chromosome are shown below. Significant differential contacts (P<0.005) between the maternal and paternal chromosome were called by 4C-ker^34^ with significant maternal contacts in red and significant paternal contacts in blue. The *Gtl2-Wdr25* interaction is highlighted in yellow. For the *Dlk1* viewpoint data (**H**) the log_2_ ratio of maternal (red) to paternal (blue) contacts are plotted below. (**I**) In maternal Δ10Ins6 heterozygote fetal brain, loss of binding at both CTCF6 and 7 causes significant reduction in expression of *Gtl2*, *Rian* and *Mirg* but no reactivation of *Dlk1*. Expression changes are independent to the destabilisation of the interaction between *Wdr25* and *Gtl2* and reduction in insulation at subTAD boundary observed in 4Cseq, which are due only to the loss of CTCF binding at CTCF7.

4Cseq analysis of the Δ10Ins6 mutants shows the same effects as in the single ΔCTCF7 line (Fig. 4G,H). The maternal chromosome-specific interaction between *Gtl2* and *Wdr25* is destabilised with no significantly different contacts called between the two chromosomes (Fig. 4G). Using the *Dlk1* promoter as the viewpoint indicates that the insulation at the Gtl2-DMR is also reduced in the Δ10Ins6 maternal heterozygotes (Fig. 4H). No conformational differences compared with wildtypes were observed in paternal Δ10Ins6 heterozygotes (Supplementary Fig. S4D and E). Together, these results suggest that CTCF binding at CTCF7 is required to stabilize the interaction between *Gtl2* and *Wdr25* and is necessary but not sufficient for the formation of the *Gtl2/Wdr25* sub-TAD boundary. Furthermore, while deletion of CTCF6 and CTCF7 alone had no influence on transcription, together both CTCF binding sites are required for the correct expression of *Gtl2, Rian* and *Mirg* in fetal brain, but this is independent of the observed conformational changes, which are also observed in the CTCF7 mutation alone.

### Maternal loss of *Wdr25* CTCFs alters conformation but not gene expression

We then turned our attention to the *Wdr25* CTCFs, which demarcate both the maternal and paternal specific proximal TAD boundaries, to investigate the role they play in allele-specific conformation and gene expression. For this, we utilized a mouse model with a 970bp deletion spanning the two CTCF binding sites (CTCF1 and 2) in intron 2 (Wdr25^em2H^ – Fig. 5A). Mice harbouring the deletion are viable and fertile, and showed no significant changes in expression levels across the domain upon maternal or paternal inheritance (Fig. 5B). No loss of imprinting was observed as indicated by the lack of significant difference in the parental expression ratio upon either maternal or paternal transmission of the deletion (Fig. 5C).

**Fig. 5.**
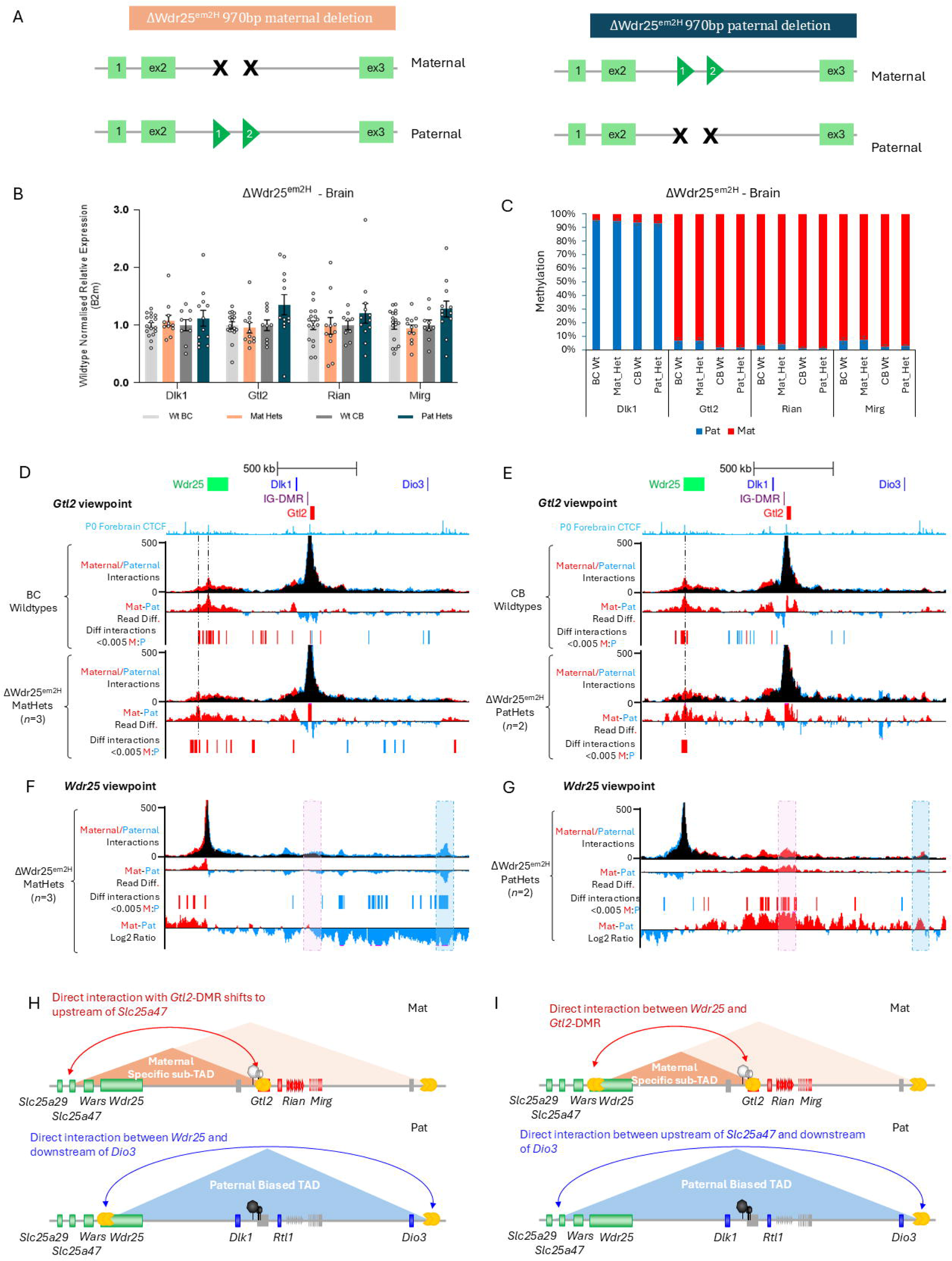
Maternal loss of Wdr25 CTCFs alters conformation but not gene expression. (**A**) Schematic of *Wdr25* intron 2 in Wdr25^em2H^ maternal and paternal heterozygotes. (**B**) Expression analysis in E15.5 brain of Wdr25^em2H^ maternal and paternal heterozygotes compared with wildtype littermates as determined by RT-qPCR. Wildtype normalised expression relative to *B2m* is shown. Wildtype BC n=17, maternal heterozygotes n=12, wildtype CB n=9, paternal heterozygotes n=11-12 Error bars represent SEM, significance of difference determined using two-tailed non-paired *t* test. (**C**) Ratio of expression from maternal (red) and paternal (blue) allele as determined by pyrosequencing in E15.5 fetal brain. Samples numbers are the same as for expression analysis. (**D**) 4Cseq data in E15.5 brain in BC wildtypes (upper panel) and Wdr25^em2H^ maternal heterozygotes (lower panel) with the *Gtl2* promoter as the viewpoint fragment. (**E**) 4Cseq data in E15.5 brain in CB wildtypes (upper panel) and Wdr25^em2H^ paternal heterozygotes (lower panel) with the *Gtl2* promoter as the viewpoint fragment. Dotted black lines indicate the highest interaction peaks. (**F and G**) 4Cseq with *Wdr25* at the viewpoint in Wdr25^em2H^ maternal heterozygotes (**F**) and paternal heterozygotes (**G**). The *Wdr25*-*Gtl2* interaction is highlighted in pink, and the *Wdr25*-*Dio3* interaction is highlighted in blue. For all 4Cseq data the mean average reads per fragment (normalised to total assigned reads) are shown. Contacts on the maternally inherited chromosome are shown in red and contacts on the paternally inherited chromosome are shown in blue. The read difference between the maternal (red) and paternal (blue) chromosome are shown. Significant differential contacts (P<0.005) between the maternal and paternal chromosome were called by 4C-ker ^34^ with significant maternal contacts in red and significant paternal contacts in blue. For the *Wdr25* viewpoint data the log_2_ ratio of maternal (red) to paternal (blue) contacts is plotted below. (**I**) In maternal Wdr25^em2H^ heterozygote fetal brains, loss both CTCF1 and 2 causes a shift to the maternal-specific *Wdr25-Gtl2* subTAD boundary to between *Slc25a29* and *Slc25a47* with no change on the paternal chromosome (**J**) In paternal Wdr25^em2H^ heterozygote fetal brains the *Wdr25-Dio3* TAD boundary to shifts to between *Slc25a29* and *Slc25a47* on the paternal chromosome, with no change seen on the maternal chromosome.

4Cseq analysis was performed on Wdr25^em2H^/CAST/EiJ heterozygotes with the *Gtl2* promoter as the viewpoint. In maternal heterozygote fetal brains, the *Gtl2/Wdr25* interaction is lost, with the peak of interaction shifting upstream so the strongest contacts are now located within *Wars* and between *Slc25a29* and *Slc25a47* (Fig. 5D and Supplementary Fig. S5A). On paternal transmission of Wdr25^em2H^ no change was seen to the *Gtl2/Wdr25* interaction on the maternally inherited chromosome, but more weak upstream contacts were observed on the paternal chromosome (Fig. 5E). 4Cseq in maternal Wdr25^em2H^ heterozygotes with *Wdr25* as the viewpoint, shows a complete loss of the interaction with *Gtl2* on the maternal chromosome (Fig. 5F), once again indicating that this conformation is not dictating expression and it is not the result of a promoter/enhancer contact. Deletion of the *Wdr25* CTCFs on the paternal chromosome, shows loss of the contact between *Wdr25* and downstream of Dio3 (Fig. 5G), consistent with the shift of the proximal TAD boundary in this model.

The shift of the TAD boundary and loss of insulation at *Wdr25* intron 2 is also evident in plots of the log₂ ratio of contact frequency between the two parental chromosomes. Both maternal and paternal heterozygotes show a clear switch in parent-of-origin contact frequency at the *Wdr25* CTCF binding site deletion. In Wdr25^em2H^ maternal heterozygotes, a higher proportion of paternal contacts are seen downstream of the *Wdr25* CTCF sites than upstream. This reflects CTCF binding at sites 1 and 2 on the paternal allele, which act as an insulator and restricts interactions beyond the TAD boundary. In contrast, on the maternally inherited chromosome, deletion of sites 1 and 2 prevents CTCF binding. This results in an upstream shift of the TAD boundary, loss of insulation and an increased proportion of maternal contacts captured upstream of the viewpoint (Fig. 5F).

The reciprocal pattern is observed in Wdr25^em2H^ paternal heterozygotes. Here, loss of CTCF binding on the paternal chromosome shifts the TAD boundary upstream and reduces insulation at the paternal *Wdr25* intron2. This causes an increased proportion of paternal contacts to be captured upstream of the viewpoint. On the maternally inherited chromosome, CTCF binds and acts to insulate so more maternal interactions captured downstream (Fig. 5G). This shift in the TAD boundary and loss of insulation has no significant impact on imprinted gene expression levels or allelic bias on either side of the TAD indicating it is not acting as a tight boundary for the imprinted domain in fetal brain (Supplementary Fig. S5B and C).

## Discussion

Here we show that the *Dlk1-Dio3* domain forms both parent-of-origin and tissue-specific chromatin conformations in vivo. In fetal brain the whole domain lies within one TAD on the paternally inherited chromosome, whilst on the maternally inherited chromosome a sub-TAD is formed between *Wdr25* and the Gtl2-DMR, a post-fertilisation acquired secondary DMR. These experiments were performed on whole fetal brain and, as brain is a heterogeneous tissue, it is possible that these findings reflect a cell-type-specific conformation, which would require further investigation. This long-range interaction is absent in fetal liver where contacts are restricted to smaller regions. The brain *Gtl2-Wdr25* sub-TAD is defined by two CTCF binding sites in intron 1 of *Gtl2* (CTCF6 and CTCF7) that are only able to bind CTCF on the unmethylated maternal chromosome^35^. Our data show these CTCF sites form a region of insulation on the maternal chromosome, restricting contacts to upstream of the Gtl2-DMR.

When the IG-DMR is deleted on the maternal chromosome in the germline, it undergoes an epigenotype switch and expresses the normally paternally expressed genes^20^. This loss of imprinting is accompanied by adoption of a paternal-chromosome-specific conformation in the fetal brain. The converse is also true when the paternal chromosome is switched to behave like a maternal chromosome by deleting the IG-CGI in the germline, it adopts a maternal-chromosome specific conformation in fetal brain^24^. This shows that parent-of-origin specific chromatin conformations observed in the developing fetus are defined by the differential epigenetic states that are established in the germline. Furthermore, tissue-specific differences in conformation between brain and liver remain upon IG-DMR deletion, indicating that tissue-specific factors act over-and-above parent-of-origin specific epigenetic modifications to establish the long-range 3D architecture in different tissues. Intriguingly, despite loss of imprinting in liver, no change in conformation is observed in maternal IG-DMR knockout mice, suggesting that, in this tissue, conformation in the domain is independent of both gene expression and germline conferred epigenetic states.

Loss of CTCF binding at CTCF7 on the maternal chromosome leads to destabilisation of the brain-specific *Gtl2-Wdr25* interaction in both the ΔCTCF7 and Δ10Ins6 models but this loss of conformation does not always occur alongside a change in expression, indicating that gene expression in the domain is independent of this conformation. This contrasts with previous data, where loss of CTCF7 binding caused reactivation of maternal *Dlk1* in ESC-derived neural stem cells^17^ and highlights the advantages of in vivo over in vitro approaches in understanding the biological relevance of particular genomic elements in specific cellular contexts. We find that in fetal brain, loss of binding at both CTCF6 and 7 in Δ10Ins6 leads to a reduction in *Gtl2* expression, suggesting that CTCF binding in intron 1 is necessary for the correct levels of *Gtl2* expression. This change in expression is independent of the loss of specific brain conformation (the same conformation change is seen in maternal ΔCTCF7s that do not change expression) and may instead be due to CTCF’s roles in recruiting either Pol II to its target sites or, as this reduction is tissue-specific, brain-specific co-factors^36^. Alternatively, CTCF6 and 7 may together act as an insulator, and the loss of CTCF binding at both sites may enable a brain-specific, downstream repressor to downregulate *Gtl2*.

Loss of CTCF binding sites 1 and 2 in *Wdr25,* which represents the 5’ boundary of the TAD, results in a change in conformation with no effect on expression or imprinting. On maternal inheritance an upstream shift of the TAD boundary occurs. This observation is not surprising as it has previously been shown that loss of CTCF binding at a boundary leads to the next adjacent upstream site being used instead and indicates the genome provides backup to maintain local conformation^9^. Deletion on the paternally inherited chromosome leads to loss of the distal *Wdr25-Dio3* interaction. No significant changes in gene expression or imprinting occur as a result of the loss of CTCF 1 and 2, once again showing that this brain-specific conformation is not required for gene regulation in the domain. Furthermore, lack of expression changes in *Wars* and *Slc25a47* when they are brought into the TAD show this boundary is not acting as a barrier for the spread to epigenetic control of the region to genes normally located outside the TAD.

The Gtl2-DMR, acquired early in development in response to the germline ICR, forms maternal chromosome-specific interactions with *Dlk1* in all tissues studied, implying it may function to repress *Dlk1* expression on the maternal chromosome. Mouse models harbouring maternal deletions of the Gtl2-DMR recapitulate the maternal IG-DMR deletion causing loss of imprinting; proportionately downregulating *Gtl2* and activating *Dlk1*^27,28^. It has been proposed that the *Gtl2* lncRNA acts to repress *Dlk1* on the maternal chromosome and, in ESCs, it has been shown to recruit PRC2 to *Dlk1*^37^. However, here we show that loss of CTCF binding in intron 1 leads to a 40% reduction in *Gtl2* expression with no effect on *Dlk1* expression. One possibility is that a 40% reduction in *Gtl2* expression is not sufficient to induce the activation of *Dlk1* expression, but a *Gtl2* truncation model has previously shown that a 20% reduction in *Gtl2* expression leads to a 33% increase *Dlk1* in expression in cardiomyocytes, suggesting this is not the case^29^. The findings here indicate that in brain *Dlk1* repression is unresponsive to this change in the *Gtl2* lncRNA, opening up the possibility that the Gtl2-DMR itself contains a direct repressor that is disrupted in the previously reported deletion models.

These findings suggest the epigenetic state of the IG-DMR, the domain ICR which is established in the germline, initiates a hierarchy of events in early development that determines both parent-of-origin specific expression and conformation (Fig. 6). This is achieved in a tissue-dependent manner as fetal brain and liver form different conformations, and loss of the ICR has no effect on conformation in the liver. Furthermore, although conformation and expression in fetal brain are both predestined by the differential methylation established in the germline, they are functionally uncoupled, suggesting that in some contexts, chromatin architecture may primarily serve a structural role in packaging DNA rather than directly facilitating gene expression.

**Fig. 6.**
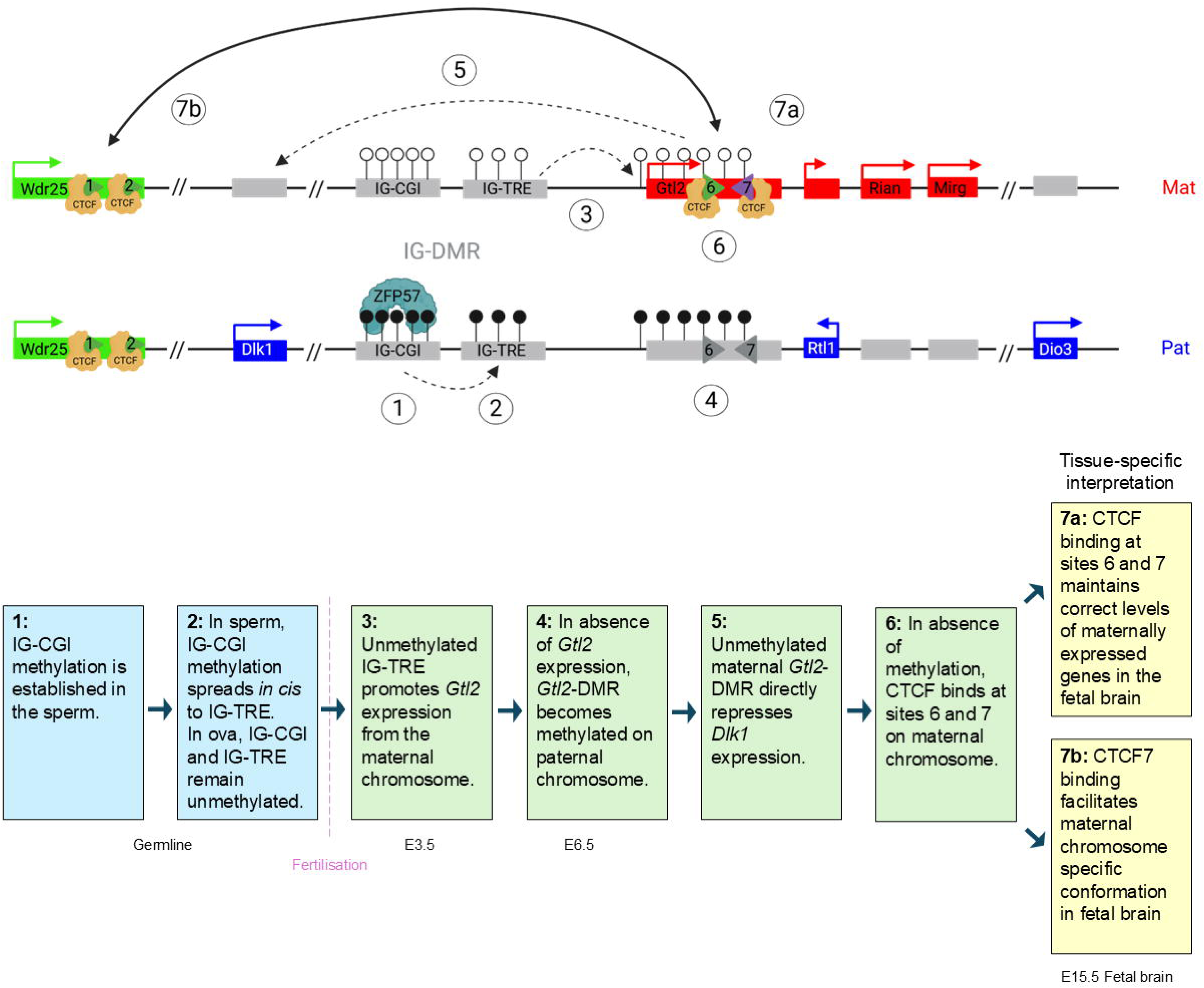
A developmental cascade independently establishes parent-of-specific expression and conformation in the fetal brain. On the maternally inherited chromosome, the IG-DMR arrives unmethylated from the ovum. The unmethylated IG-TRE acts as an enhancer causing *Gtl2* to be monoallelically expressed and the Gtl2-DMR region to remain unmethylated on the maternal chromosome. The unmethylated Gtl2-DMR can then directly repress *Dlk1* expression independently of the ncRNA expression. CTCF binds unmethylated sites 6 and 7 in fetal brain, this monoallelic CTCF binding independently influences the expression levels of all the maternally expressed genes. Monoallelic binding of CTCF at site 7 facilitates the maternal-chromosome specific interaction with *Wdr25*. On the paternally inherited chromosome, the IG-CGI and IG-TRE become methylated in sperm. Methylation at the IG-TRE means it is unable to enhance *Gtl2* expression and this in turn leads to the *Gtl2* promoter and intron 1 becoming methylated. Methylation of the Gtl2-DMR region allows *Dlk1* expression on the paternal chromosome. It also prevents CTCF from binding to sites 6 and 7 and instead allows for the CTCF binding sites in Wdr25 to interact with those downstream of *Dio3* forming the paternal chromosome specific conformation in fetal brain. Adapted from Weinberg-Shukron et al 2023 ^38^

## Materials and Methods

### Mice

All animal experiments were performed in accordance with the Animals (Scientific Procedures) Act 1986 Amendment Regulations 2012 following ethical review by the University of Cambridge Animal Welfare and Ethical Review Body. Animal experiments were approved by the UK Home Office project license #PP8193772. All efforts were made to minimize animal discomfort. For the hybrid analysis with *Gtl2*, *Dlk1* and *Wdr25* as viewpoints, reciprocal crosses of the mouse lines C57BL/6J and castaneus (CAST/EiJ) were generated. For the loss of IG-DMR, IG-CGI and CTCF6 and 7 studies, female or male heterozygotes were mated to Mus castaneus (CAST/EiJ) mice to generate heterozygotic conceptuses and wildtype littermates.

### Generation of *Gtl2* CTCF deletion mice

The gRNAs were designed using the CRISPR MIT tool from the Zhang lab (crispr.mit.edu). The guides were as follows:

CTCF6_1F: 5’-TAATACGACTCACTATAGTTCTATTGATGGCCGC-3’

CTCF6_1R: 5’-TTCTAGCTCTAAAACCTTGCGGCCATCAATAGAA-3’

CTCF7_2F: 5’-TAATACGACTCACTATAGGCACATAGAGACCGCT-3’

CTCF7_2R: 5’-TTCTAGCTCTAAAACCCTAGCGGTCTCTATGTGC-3’

They were synthesised following the GeneArt Precision gRNA kit protocol. 50ng/μL of gRNA were mixed with 42ng/μL Cas9 protein (Gene Art Platinum Cas9 nuclease). Superovulated female (C57Bl6JxCBA)F1 mice were mated to (C57Bl6JxCBA)F1 stud males. The gRNA and Cas9 mRNA/protein mix was injected into fertilized eggs, and 15–25 blastocysts were transferred into uterus of pseudopregnant females. The Δ10Ins6 mice were generated by simultaneously injecting both guides into zygotes. The zygote injection was performed at the Cambridge Stem Cell Facility.

Wdr25^em2H^ mice were generated as part of the Genome Editing Mice for Medicine programme (GEMM) at the Mary Lyon Centre.

### Genotyping

For genotyping, DNA was extracted using PCR Biosystems PCRBIO Rapid extract PCR kit (PB10.24-40), following manufacturer’s instructions. PCRs were performed using either KOD Hot Start DNA polymerase (Sigma-Aldrich #71086-3) or REDTaq® ReadyMix™ PCR Reaction Mix (Sigma-Aldrich #R2523) following manufacturer’s instructions and cycling conditions listed in Supplementary Table S1. Genotyping PCR products were Sanger sequenced to characterise the deletions. Mouse lines were backcrossed to C57BL/6J for at least 10 generations to minimise any off-target effects.

### 4Cseq

The 4Cseq protocol is based on previously published methodologies^32,33^. Whole brain and liver were harvested from E15.5 (15.5 days after conception). Brains were homogenised and digested with 200U/ml collagenase (type IV Gibco™17104-019) in 250μl PBS+10% FBS and incubated at 37°C for 45 minutes. Cells from brains and non-collagenase treated livers were the dissociated through 70μm sieve. Samples were then centrifuged at 400g for 8 minutes at 4°C, the supernatant removed, and cells resuspended in 5ml PBS/10%FBS. Cells were then fixed in 4% formaldehyde in PBS for 10 minutes with rolling. The fixation reaction was quenched with glycine (final concentration 0.125M) on ice. Samples were centrifuged at 400g for 8 min, at 4°C, washed with cold PBS, then centrifuged again and the supernatant removed. Samples were resuspended in 1ml cold lysis buffer (10 mM Tris–HCl pH 8, 10 mM NaCl, 0.2% Igepal (CA-630 Sigma) and 1x cOmplete™, EDTA-free Protease Inhibitor Cocktail (Roche # 11873580001)) and incubated on ice for 30 min with occasional mixing. Nuclei were centrifuged at 750g for 5 min at 4°C and resuspended in 1x PBS. 5x10^6^ nuclei aliquots were pelleted at 800g at 4°C for 5 min, supernatant removed then flash frozen. Pellets were stored at -70°C.

The nuclei were resuspended in 1.1x restriction enzyme buffer and 2.86% SDS and incubated at 37°C 1 hr with shaking at 900 rpm. TX-100 was added to a final concentration of 1x buffer, 2.5%SDS and 1.25% TX-100 then incubated at 37°C 1 hr with shaking at 900 rpm. 600U of the primary cutter (DpnII) was added in three steps of 200U over a 24-hr incubation at 37°C while shaking at 900 RPM, then heat inactivated at 65°C for 20 min. Samples were ligated with 50U T4 DNA ligase in a 7ml reaction overnight at 16°C. Cross-links were reversed by adding 300μg proteinase K and incubated overnight at 65°C. 300μg of RNase A were added and incubated at 37°C for 45 min. Samples were then Phenol-Chloroform extracted and EtOH precipitated.

Samples were digested with 50U of the secondary cutter (NlaIII) in a 500ul reaction and incubated overnight at 37°C, then heat inactivated at 65°C for 25min. The second ligation was performed in a 14ml reaction with 100U of T4 ligase incubated overnight at 16°C. Samples were EtOH precipitated overnight, resuspended in 150ul 10mM Tris-HCl pH7.5, then purified using QIAquick PCR purification kit (Qiagen #28104). 3.2μg of 4C template were then amplified with viewpoint specific primers (Supplementary Table S1) in 16 50μl reactions using primer specific PCR conditions (Supplementary Table 1). The reactions were then pooled, purified using the High Pure PCR Product Purification Kit (Roche) eluting in DDW. Libraries were size selected using BluePippin (BDF2003 2% agarose gel cassette. Dye-free with internal standards) or Ampure beads (Beckman). Libraries were sequenced as detailed in (Supplementary Table S1).

### Data Processing

For the *Gtl2* viewpoint in C57BL/6xCAST/EiJ reciprocal hybrids, paired end reads were first separated into C57BL/6 or CAST/Eij using a SNP in read 2 using Cutadapt^39^. The libraries were then separated using the 3bp barcode and the forward library primer and any adapter sequence was removed using trim_galore (v0.4.1) (Babraham Bioinformatics - Trim Galore!). For the *Dlk1* and *Wdr25* viewpoint in C57BL/6xCAST/Eij reciprocal hybrids libraries were first separated using the 3bp barcode in the forward library primer, then they were assigned to either C57BL/6 or CAST/Eij using a SNP in read 1 before any adapter sequence was removed using trim_galore (v0.4.1). Reads from 150 bp libraries were trimmed to 50 bp to remove any chimeric interactions and aid alignment.

C57BL/6 reads were aligned to GRCm38(mm10) and CAST/Eij reads were aligned to a Mus Castaneus mm10 pseudo-genome using Bowtie2^40^. Alignments were converted to bam files using SAMtools^41^. The 4Cseq aligned libraries were analysed using the Basic4C-seq package in R^42^. The C57BL/6 and CAST/Eij libraries were aligned to specific C57BL/6 or CAST/Eij “virtual fragment libraries” generated with the particular first and second cutters used to generate the libraries. Near cis analysis was performed on the region spanned by three previously reported TADs (chr12:105801790-113201789) and the counts normalised to library size^1^. A read distribution analysis was performed in Basic4C-seq to assess library quality. The mean of the “fragEndReadsAverage” across all biological replicates was calculated and converted to a bigwig file to display as a track hub on UCSC genome browser^31^. The read difference between the maternal and paternal contacts was then calculated. For the log_2_ tracks, the ratio between the mean maternal “fragEndReadsAverage” +1 and mean paternal “fragEndReadsAverage” +1 was calculated and log_2_ transformed. For the read difference and log_2_ tracks, only the fragments with the same coordinates between the C57BL/6 and CAST/Eij genomes were used.

### Statistical analysis

All 4Cseq libraries were generated in hybrid mice; this allows the two parentally inherited chromosomes to be distinguished from each other using SNPs. Hybrid libraries also control for the number of cells as the reads from each chromosome arise from the same starting material. This allowed significantly differential contacts between the maternal and paternal chromosome to be called using 4C-ker^34^. 4C-ker uses Hidden Markov Models to partition the genome and DEseq2^43^ to call differential interactions. A previous benchmarking paper has shown this to be the most sensitive algorithm for calling interaction differences between conditions^44^. Parent-of-origin specific interactions were called if statistically different contacts (with the P value set at <0.005) were identified between the two chromosomes in wildtypes from both reciprocal crosses. Lack of significant differences between the two chromosomes in mice that are heterozygotic for a CTCF deletion indicate loss of parent-of-origin specific topologies.

### Reverse transcription and quantitative real-time PCR (qPCR)

Total RNA was isolated from E15.5 brain or liver using Tri Reagent (Sigma-Aldrich #T9424) according to the manufacturer’s instructions, quantified by spectrophotometry and quality controlled by gel electrophoresis to assess the integrity of the ribosomal RNA and absence of degradation products. RNA was treated with RNase-free DNase I (Thermo Fisher Scientific #EN0521) prior to cDNA synthesis using RevertAid H Minus First Strand cDNA Synthesis Kit with random hexamer primers (Thermo Fisher Scientific #K1632). qPCR was performed with Brilliant II SYBR® Green QPCR Master Mix (Agilent Technologies #600882) in a 384-well plate on a QuantStudio™ 5 Real-Time PCR System (Applied Biosystems #A34322) using primers listed in Supplementary Table S3. Relative quantification of gene expression was normalized to beta-2 microglobulin expression levels and calculated using the ΔCT method. The relative expression levels were normalised to the mean of the wildtype littermates and the mean normalised values across all litters are plotted. A two-tailed non-paired *t* test was used to access the significance of any change in expression levels compared with wildtype litter mates.

### Allele-Specific pyrosequencing

Pyrosequencing was carried on a PSQ HS96 System using PyroMark Gold Q96 SQA Reagents (Qiagen #972812) and products of the qPCR detailed above. SNP percentage was determined by the pyromarkTM MD 1.0 software. All sequencing primers used are listed in Supplementary Table S3. A two-tailed non-paired *t* test was used to access the significance of any change in allelic ratio levels compared with wildtype litter mates.

### Chromatin immunoprecipitation

ChIP was performed as previously described with some modifications^45,46^. Primary niche astrocytes were cultured from the subventricular zone (SVZ) harvested from the CTCF mutant mice and wildtype littermates at P0. SVZ tissue was transferred into 1ml of HBSS (company), dissociated and transferred into a solution of 5ml of 0.25% trypsin and 100units of DNaseI. The samples were shaken at room temperature for 30 minutes, centrifuged at 1500rpm for 5 minutes and the supernatant removed. The pellet was dissociated twice in 1ml of NB/FBS then centrifuged at 2000rpm for 5 minutes, supernatant was removed, and cells were resuspended in 1ml of NB/FBS. The dissociated single cells were transferred into a preprepared T25 flask (coated in Poly-D-Lysine (PDL) for at least 1 hour and washed three times with molecular water) with a total of 5ml of NB/FBS medium (NuroBasal supplemented with 5% certified FBS, 3% Pen/Strep, 2% B27, 1% Glutamax) to enrich for neuronal populations. Astrocytes were enriched for by removing other neuronal populations by washing vigorously in PBS the day after seeding and before splitting the culture. Cells were incubated at 37°C and media was replaced every 2-3 day and after ∼7 days the cells were split into T75 flasks and incubated at 37°C for a further 7 days with media changes every 2-3 days before collection. 1x10^7^ primary niche astrocytes were resuspended in 5ml PBS/10%FBS and crosslinked as described in the 4Cseq protocol above.

The nuclei were thawed on ice, resuspended in SDS shearing lysis buffer (10mM Tris-HCl pH 8, 1mM EDTA, 0.15% SDS and EDTA-free cOmplete™), and sonicated (Bioruptor Plus, Diagenode) at 4°C to generate DNA fragments of 100-500bp (3 repetitions of 10 sonication cycles of 30 seconds on and 30 seconds off). The sonicated lysates were subsequently clarified by centrifugation (15,000 rpm for 10 minutes at 4°C). 10% of the mixture was taken as input control. The sonicated lysate was topped up to 1mL with LB3-500NaCl (20mM Tris-HCl, pH 7.5, 500mM NaCl, 1mM EDTA, 0.5mM EGTA, 1% Triton-X-100, 0.1% Sodium deoxycholate, 0.1% SDS and EDTA-free cOmplete™). This mixture was incubated overnight at 4°C with beads (Protein A Dynabeads, Invitrogen #10002D) that had been pre-blocked with 0.5% BSA and mixed with 3.3μg Rabbit polyclonal CTCF antibody (Diagenode #C15410210-50). To remove non-specifically bound proteins from the ChIP beads were washed five times with RIPA Buffer (50mM HEPES-KOH, pH 7.4, 500mM LiCl, 1mM EDTA, 1% NP-40 and 0.7% Sodium deoxycholate) and once with TE Buffer (50mM Tris-HCl, pH 8.0, 10mM EDTA) at 4°C. The DNA-protein complex was eluted from the beads in Elution Buffer (50mM Tris-HCl, pH 8.0, 10mM EDTA and 1% SDS) for 20 minutes at 65°C, and reverse-crosslinked overnight at 65°C. The eluted samples were then treated with 10mg/ml RNase A (Sigma-Aldrich #R4875) and 20mg/ml Proteinase K and purified using the QIAquick PCR Purification Kit.

qPCR was performed with Brilliant II SYBR® Green qPCR Master Mix (Agilent Technologies cat#600882) in a 384-well plate on a QuantStudio™ 5 Real-Time PCR System (Applied Biosystems cat#A34322). The ChIP-qPCR fold enrichments were calculated according to the percent input method, where signals obtained from the ChIP are divided by signals obtained from the input sample. qPCR primers are listed in Supplementary Table S5.

## Author contributions

C.A.E and A.C.F-S conceived the project, designed experiments. F.L.D, AW-S, A.M-B and D.J.G performed the experiments. F.L.D and A.K.F maintained mouse colonies. C.A.E analysed data from all experiments and prepared the manuscript.

## Acknowledgements

We thank the Ferguson-Smith group members for their discussion and advice. The Wdr25^em2H^ mice used in this study were generated as part of the GEMM programme at the Mary Lyon Centre at MRC Harwell and the following award is acknowledged: MC_UP_2201/2. Fig. 6 was created with BioRender.com.

## Data Availability

Data generated in this study will be available after publication.

