## Supplementary information for "Genomic conformation reflects epigenetic programming rather than gene expression"

**Supplementary Figure Legends**

**Supplementary Fig. S1.**

**(A**) Relative expression levels of *Dlk1* and *Gtl2* in E15.5 wildtype brain and liver from C57BL/6Jx CAST/EiJ (BC) hybrid mice. Expression levels are shown relative to B2m expression, error bars represent SEM. **(B**) 4Cseq data in E15.5 fetal brain (upper panel) and fetal liver (lower panel) with the *Gtl2* promoter as the viewpoint fragment in CAST/EiJ x C57BL/6 (CB) hybrid mice. (**C**) 4Cseq data with *Wdr25* exon 2 as the viewpoint fragment in E15.5 CB hybrid fetal brain (**D**) 4Cseq data with *Dlk1* promoter as the viewpoint in E15.5 CB hybrid fetal brain. (**E**) Zoom in on “switch-point” at Gtl2-DMR. For all 4Cseq data the mean average reads per fragment (normalised to total assigned reads) are shown. Contacts on the maternally inherited chromosome are shown in red and contacts on the paternally inherited chromosome are shown in blue. For *Gtl2* (**B**) and *Wdr25* (**C**) viewpoints the read difference between the maternal (red) and paternal (blue) chromosome are shown below. For the *Dlk1* viewpoint data (**D,E**) the log_2_ ratio of maternal (red) to paternal (blue) contacts are plotted below. Forebrain and e14.5 liver CTCF (turquoise) and E15.5 forebrain H3K27ac (Yellow) ChIP data are taken from ENCODE track hub ^30^. (**F-G**) Zoom in on the *Wdr25* interaction with Gtl2 viewpoint (**F**) and the *Gtl2* interaction with *Wdr25* viewpoint (**G**). Below, conservation of CTCF1 & CTCF2 (**F**) and CTCF6 & CTCF7 (**G**) in mammals. Contacts on the maternal chromosome are shown in red, contacts on the paternal chromosome in blue and overlapping contacts are shown in black. Motif conservation was taken from MultiZ track in UCSC ^47^ All data are viewed on the UCSC genome browser ^31^.

**Supplementary Fig. S2.**

(**A-C**) 4Cseq in fetal brain of IG-DMR deletion paternal heterozygotes (Paternal ΔIG) for the *Gtl2* viewpoint (**A**), *Wdr25* viewpoint (**B**) and *Dlk1* viewpoint (**C**). For 4Cseq data, the mean average reads per fragment (normalised to total assigned reads) are shown. Contacts on the maternal chromosome are shown in red and contacts on the paternal chromosome in blue. For *Gtl2* and *Wdr25* viewpoints the read difference between the maternal (red) and paternal (blue) chromosome are shown below. Significant differential contacts (P<0.005) between the maternal and paternal chromosome were called by 4C-ker^34^ for the *Gtl2* (**A**) and *Wdr25* (**B**). significant maternal contacts are shown in red and significant paternal contacts are shown in blue. For the Dlk1 viewpoint data (**C**) the log_2_ ratio of maternal (red) to paternal (blue) contacts are plotted below. (**D**) The paternally inherited chromosome maintains conformation up deletion of the IG-DMR. (**E**) 4Cseq in fetal liver of IG-DMR deletion maternal heterozygotes (Maternal ΔIG) for the *Gtl2* viewpoint.

**Supplementary Fig. S3.1.**

(A) Details of the three Gtl2-DMR CTCF mutation lines generated. (B) UCSC screen shot showing position of mutations relative to CTCF binding peaks ^31^. (C) CTCF ChIP in niche astrocytes derived from P0 brains of maternal and paternal ΔCTCF6, ΔCTCF7 and Δ10Ins6 heterozygotes and their wildtype littermates. Fold enrichments were calculated according to the percent input method, where signals obtained from the ChIP are divided by signals obtained from the input sample. Error bars represent SEMs. Perturbation of CTCF binding was only seen on maternal inheritance of the various mutations.

**Supplementary Fig. S3.2.**

**(**A) Schematic of the Gtl2-DMR in ΔCTCF6 and ΔCTCF7 paternal heterozygotes. (**B**) Expression analysis of ΔCTCF6 and ΔCTCF7 paternal heterozygotes compared with wildtype littermates (WT CBs) as determined by RT-qPCR. Wildtype normalised expression relative to *B2m* is shown. Error bars represent SEM, significance of difference determined using two-tailed non-paired *t* test. ΔCTCF6 WT CB n = 11, ΔCTCF6 Pat Het n = 9, ΔCTCF7 WT CB n = 15, ΔCTCF7 Pat Het n = 12, (**C**) Ratio of expression from maternal (red) and paternal (blue) allele as determined by pyrosequencing. Error bars represent standard deviation; sample numbers are the same as for expression analysis. Significance of difference determined using two-tailed non-paired *t* test.(**D-E**) 4Cseq in fetal brains of ΔCTCF6 and ΔCTCF7 paternal heterozygotes for the *Gtl2* viewpoint (**D**), and *Dlk1* viewpoint (**E**). For 4Cseq data, the mean average reads per fragment (normalised to total assigned reads) are shown. Contacts on the maternal chromosome are shown in red, contacts on the paternal chromosome in blue and overlapping contact are shown in black. For the *Gtl2* viewpoint (**D**), the read difference between the maternal (red) and paternal (blue) chromosome are shown below. Significant differential contacts (P<0.005) between the maternal and paternal chromosome were called by 4C-ker ^34^ with significant maternal contacts in red and significant paternal contacts in blue the *Gtl2-Wdr25* interaction is highlighted in yellow. For the *Dlk1* viewpoint data (**E**) the log_2_ ratio of maternal (red) to paternal (blue) contacts are plotted below.

**Supplementary Fig. S3.3.**

(**A-B**) Expression analysis of *Dlk1-Dio3* domain in E15.5 brain of ΔCTCF6, ΔCTCF7 and Δ10Ins6 maternal heterozygotes (**A**) and paternal heterozygotes (**B**) compared with their wildtype littermates as determined by RT-qPCR. Expression relative to the housekeeping gene, *B2m*, is shown. Error bars represent SEM. (**C-D**) Proportion of expression from maternal (red) and paternal (blue) allele in E15.5 fetal brain in ΔCTCF6, ΔCTCF7 and Δ10Ins6 maternal heterozygotes (**C**) and paternal heterozygotes (**D**). The ratio determined by pyrosequencing is overlaid onto wildtype normalised *B2m* relative expression levels. Error bars show SEM of expression levels.

**Supplementary Fig. S4.**

(**A**) Schematic of the Gtl2-DMR in Δ10Ins6 paternal heterozygotes. (**B**) Expression analysis in E15.5 brain of Δ10Ins6 paternal heterozygotes compared with wildtype littermates as determined by RT-qPCR. Wildtype normalised expression relative to *B2m* is shown. Wildtype n=13, heterozygotes n=19. Error bars represent SEM, significance of difference determined using two-tailed non-paired *t* test. (**C**) Ratio of expression from maternal (red) and paternal (blue) allele as determined by pyrosequencing in E15.5 fetal brain. Error bars represent standard deviation; sample numbers are the same as for expression analysis. (**D-E**) 4Cseq in E15.5 fetal brains of Δ10Ins6 paternal heterozygotes for the *Gtl2* viewpoint (**D**), and *Dlk1* viewpoint (**E**). For 4Cseq data, the mean average reads per fragment (normalised to total assigned reads) are shown. Contacts on the maternal chromosome are shown in red, contacts on the paternal chromosome in blue and overlapping contact are shown in black. For the Gtl2 viewpoint (**D**), the read difference between the maternal (red) and paternal (blue) chromosome are shown below. Significant differential contacts (P<0.005) between the maternal and paternal chromosome were called by 4C-ker ^34^ with significant maternal contacts in red and significant paternal contacts in blue. The *Gtl2-Wdr25* interaction is highlighted in yellow. For the *Dlk1* viewpoint data (**E**) the log_2_ ratio of maternal (red) to paternal (blue) contacts are plotted below

**Supplementary Fig. S5.**

**(A)** Zoom in on 4Cseq data in BC wildtypes and Wdr25^em2H^ maternal heterozygotes, interaction peaks are shown by black dotted lines, (**B**) Expression analysis of genes at the periphery of the *Dlk1-Dio3* domain in E15.5 brain of Wdr25^em2H^ maternal and paternal heterozygotes compared with wildtype littermates as determined by RT-qPCR. Wildtype normalised expression relative to *B2m* is shown. Wildtype BC n=17, maternal heterozygotes n=11-12, wildtype CB n=9, paternal heterozygotes n=11-12. Error bars represent SEM, significance of difference determined using two-tailed non-paired *t* test. (**C**) Ratio of expression of peripheral genes from maternal (red) and paternal (blue) allele as determined by pyrosequencing in E15.5 fetal brain. Sample numbers are the same as expression analysis.

**Supplemental Tables**

Supplementary Table S1 – Genotyping primers and PCR conditions

| **Model** | **Forward primer** | **Reverse primer** | **PCR conditions** |
| --- | --- | --- | --- |
| ΔCTCF6 | 5'-GCGACGTGGCAGGTTTCTAG | 5'-GTCCCGCAACCCATCTCG | KOD |
| ΔCTCF7 | 5'-AGGTGGTTGGGCTATTGGAG | 5'-GCAATGGGAGGGGTACAGA | red taq |
| Δ10Ins6 | 5'-GCGACGTGGCAGGTTTCTAG | 5'-GTCCCGCAACCCATCTCG | KOD |
| CTCFWdrKO | 5'-ACCCCATAAAGACCTGCCTC | 5'-TGGCTCCTCACATCAGTCAG | red taq |
| CTCFWdrWT | 5'-ACCCCATAAAGACCTGCCTC | 5'-ATCCCACACACAGCCCTAAA | red taq |
|  | **Cycling conditions with red taq:** | **Cycling conditions with KOD** |  |
|  | 94^o^C for 2 mins | 95^o^C 2mins |  |
|  | 94^o^C 30sec | 95^o^C 20sec |  |
|  | annealing temp ^o^C 30sec | annealing temp ^o^C 10sec |  |
|  | 72^o^C 30sec | 70^o^C 30sec |  |
|  | from step 2 for additional 34 cycles | from step 2 additional 36 cycles |  |
|  | 72^o^C for 5mins |  |  |

Supplementary Table S2 – 4Cseq assays, primers and cycling conditions.

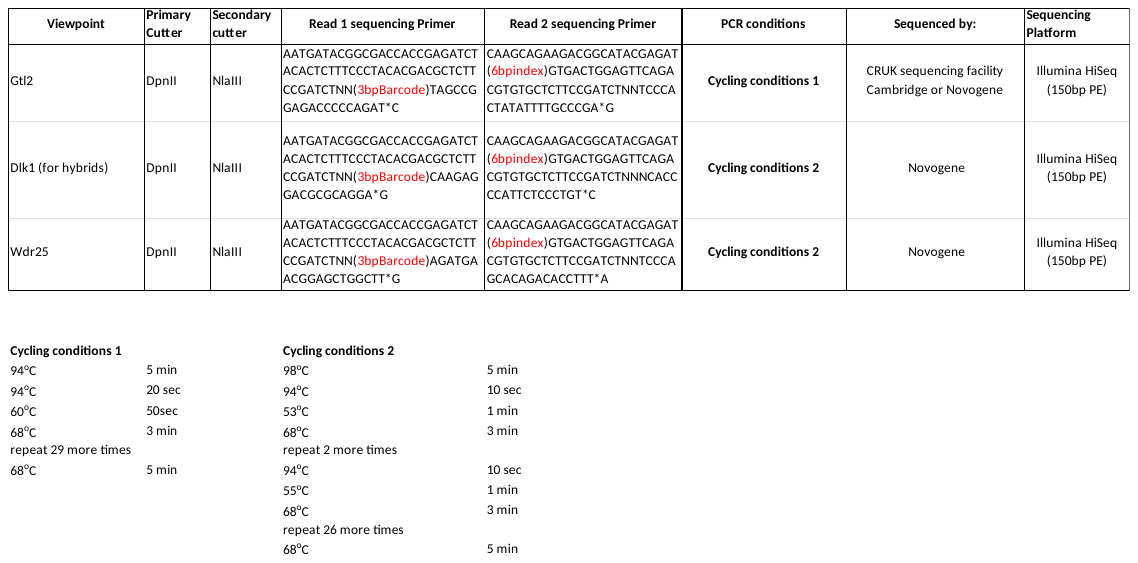

Supplementary Table S3 – qRT-PCR and pyrosequencing primers for expression analysis

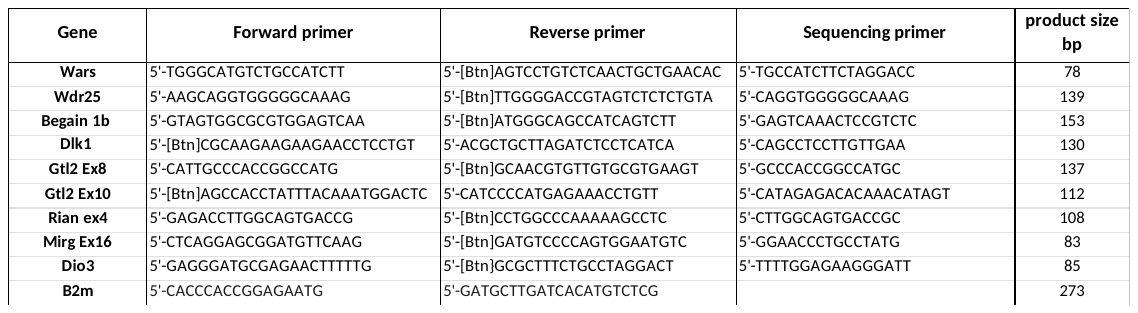

Supplementary Table S4 – Bisulfite pyrosequencing primers for methylation analysis

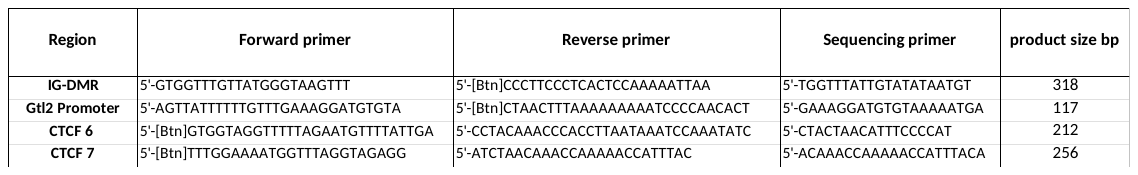

Supplementary Table S5 – Primers for ChIP qPCR analysis

| **Region** | **Forward primer** | **Reverse primer** |
| --- | --- | --- |
| CTCF6 | 5'-TCCTCATTCTGGGCTGGC | 5'-AAAATTCCGCAGTGCCACC |
| CTCF7 | 5'-CCCCCTCCTACGTCAGTCTA | 5'-AGGTCACAAGTGTTAGCTGTGTG |
| H19_DMD | 5'-CTCTGCAAAGAATCCTTTGTGTG | 5'-GCGGTGGCAGCATACTCCTATAT |
