## Supplementary figures and images for "Genomic conformation reflects epigenetic programming rather than gene expression"

### Supplementary Figure 1

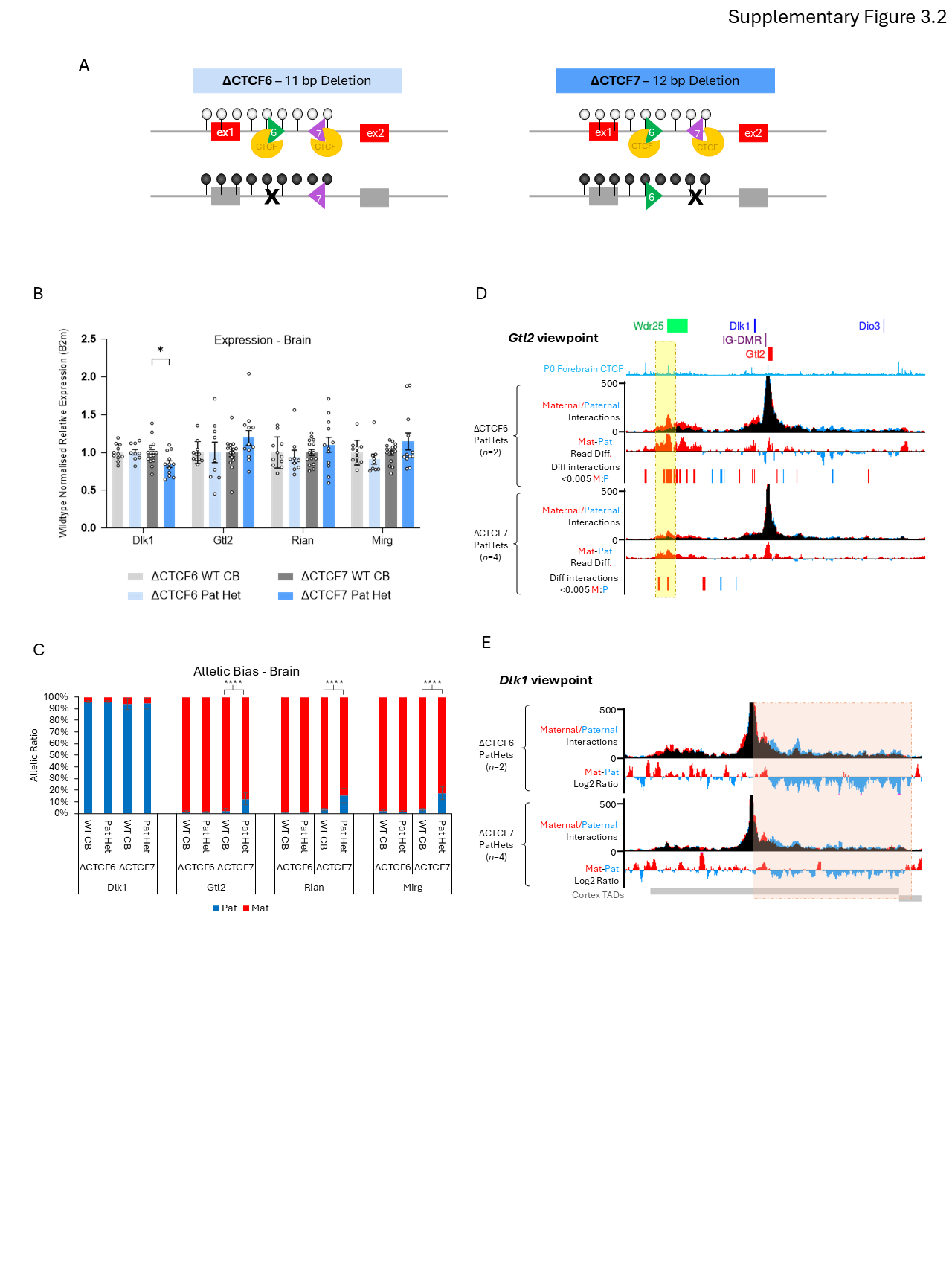

### Supplementary Figure 2

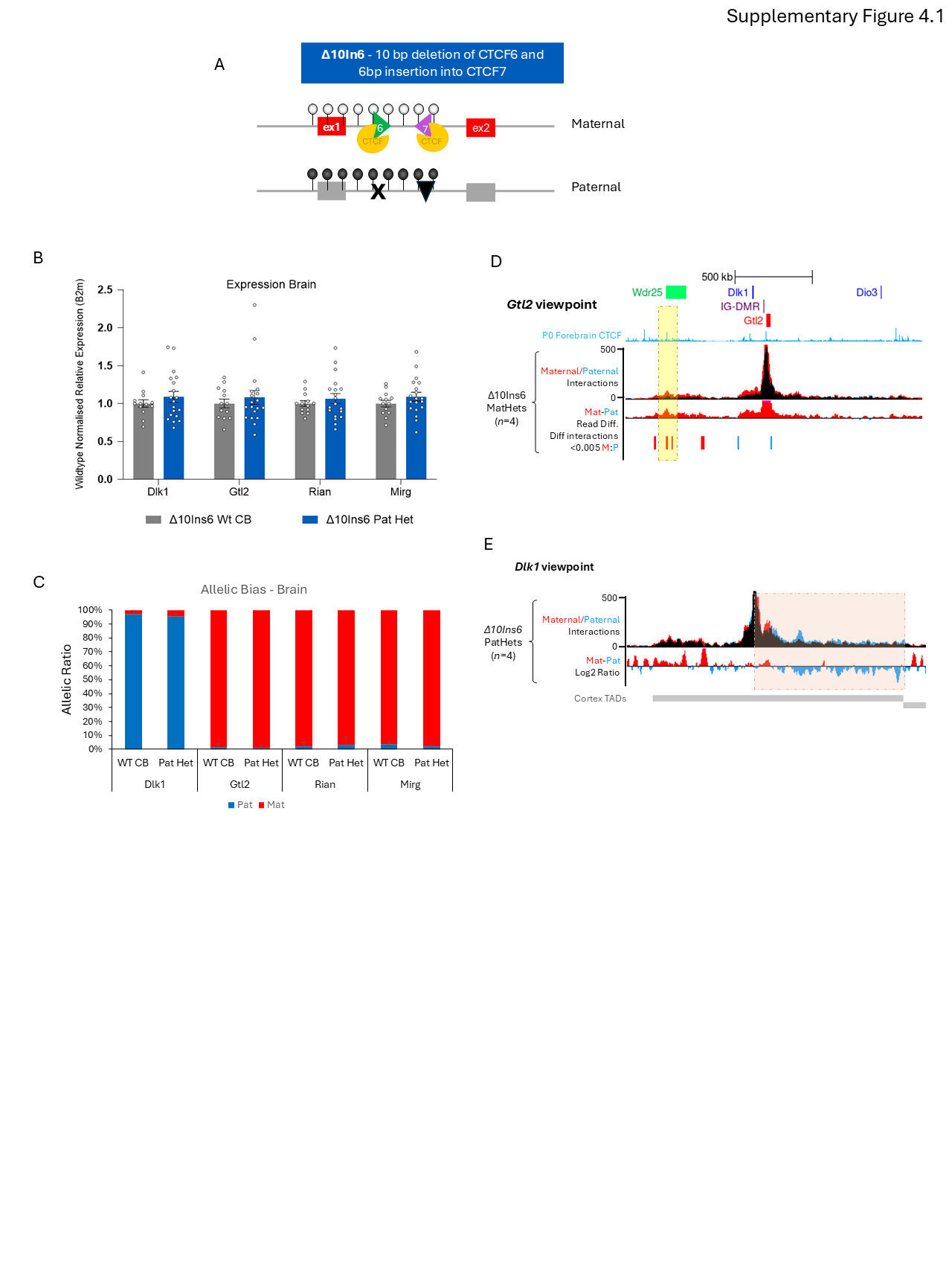

### Supplementary Figure 3.1

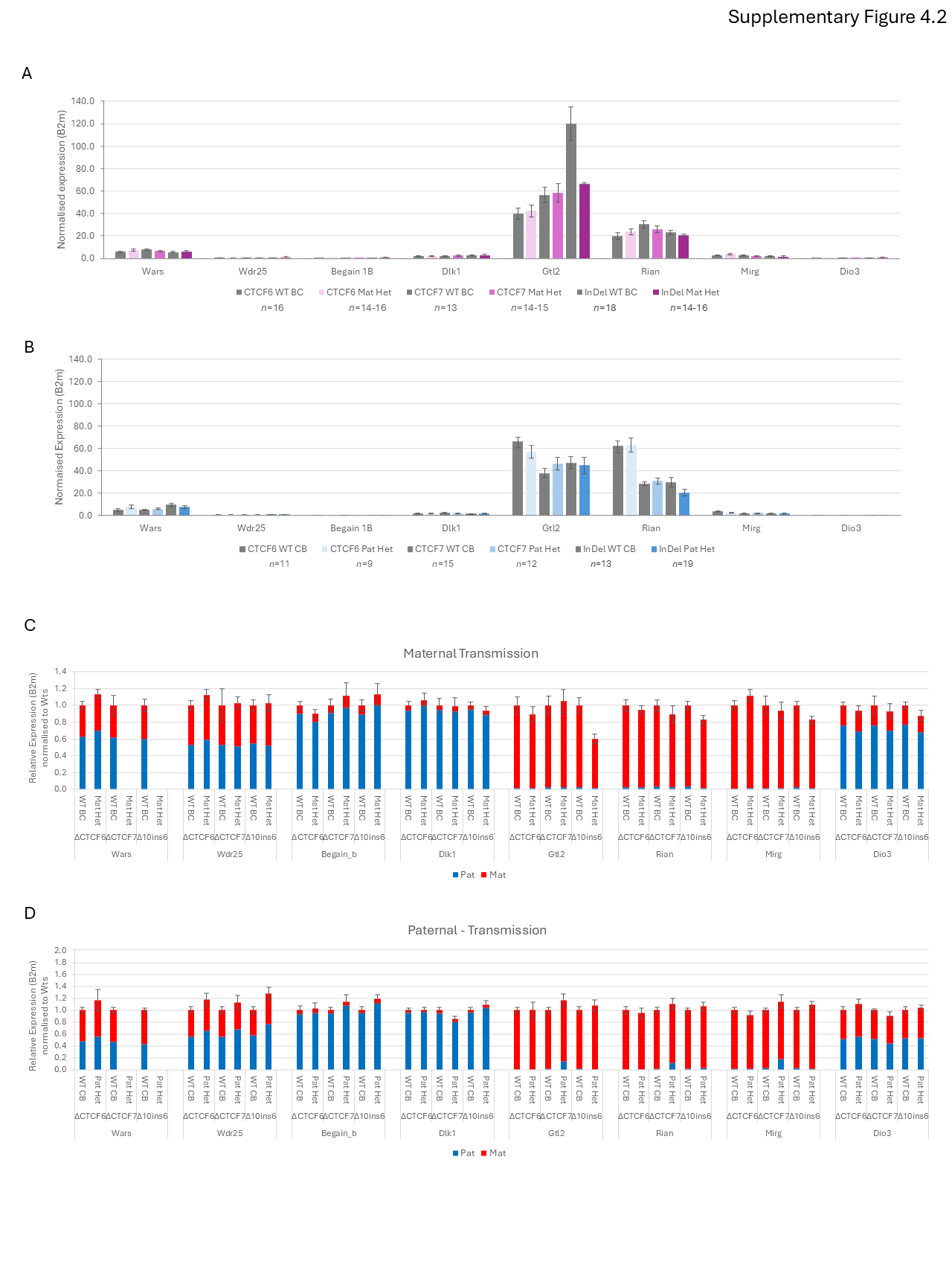

### Supplementary Figure 3.2

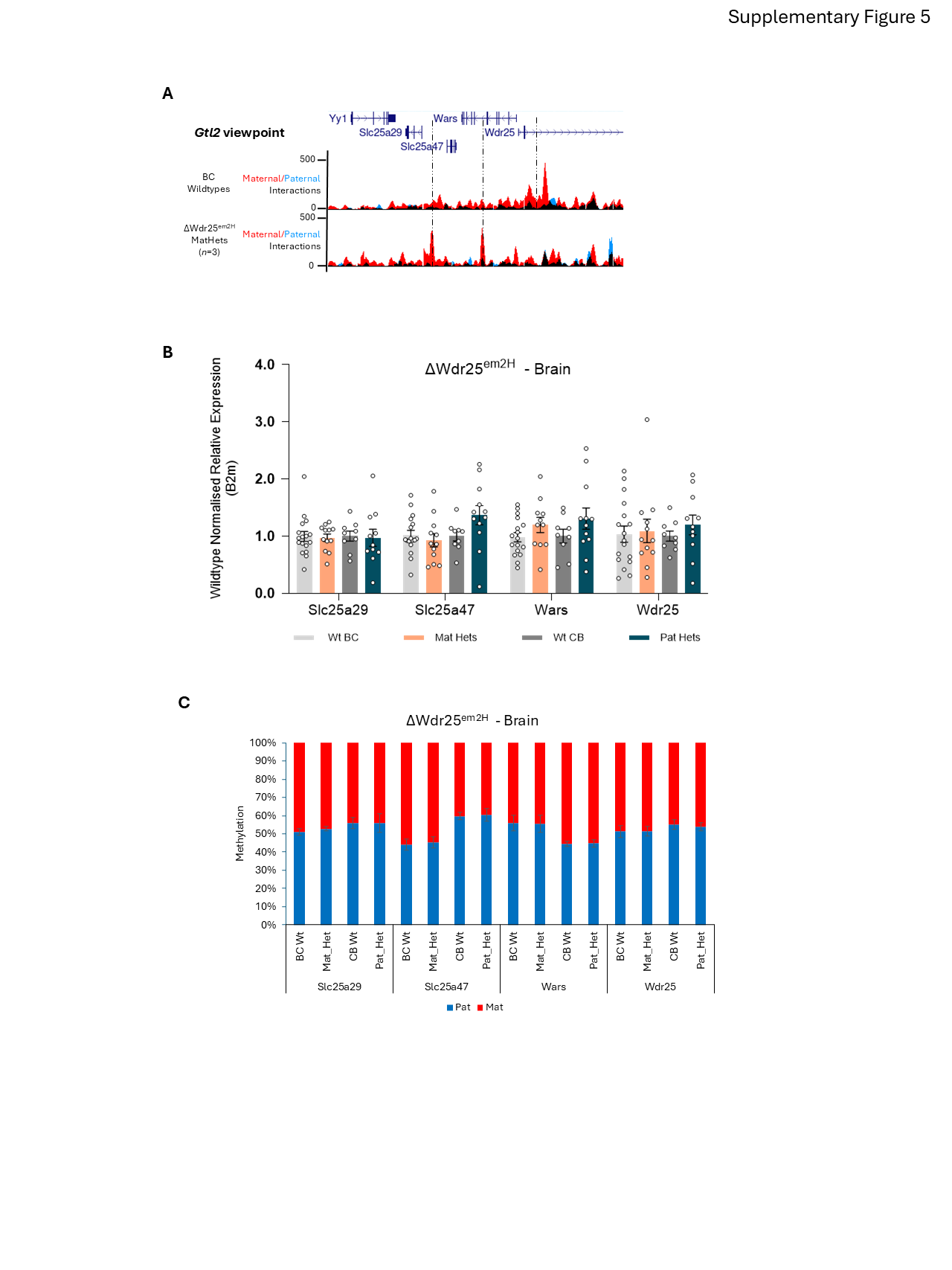

### Supplementary Figure 4.1

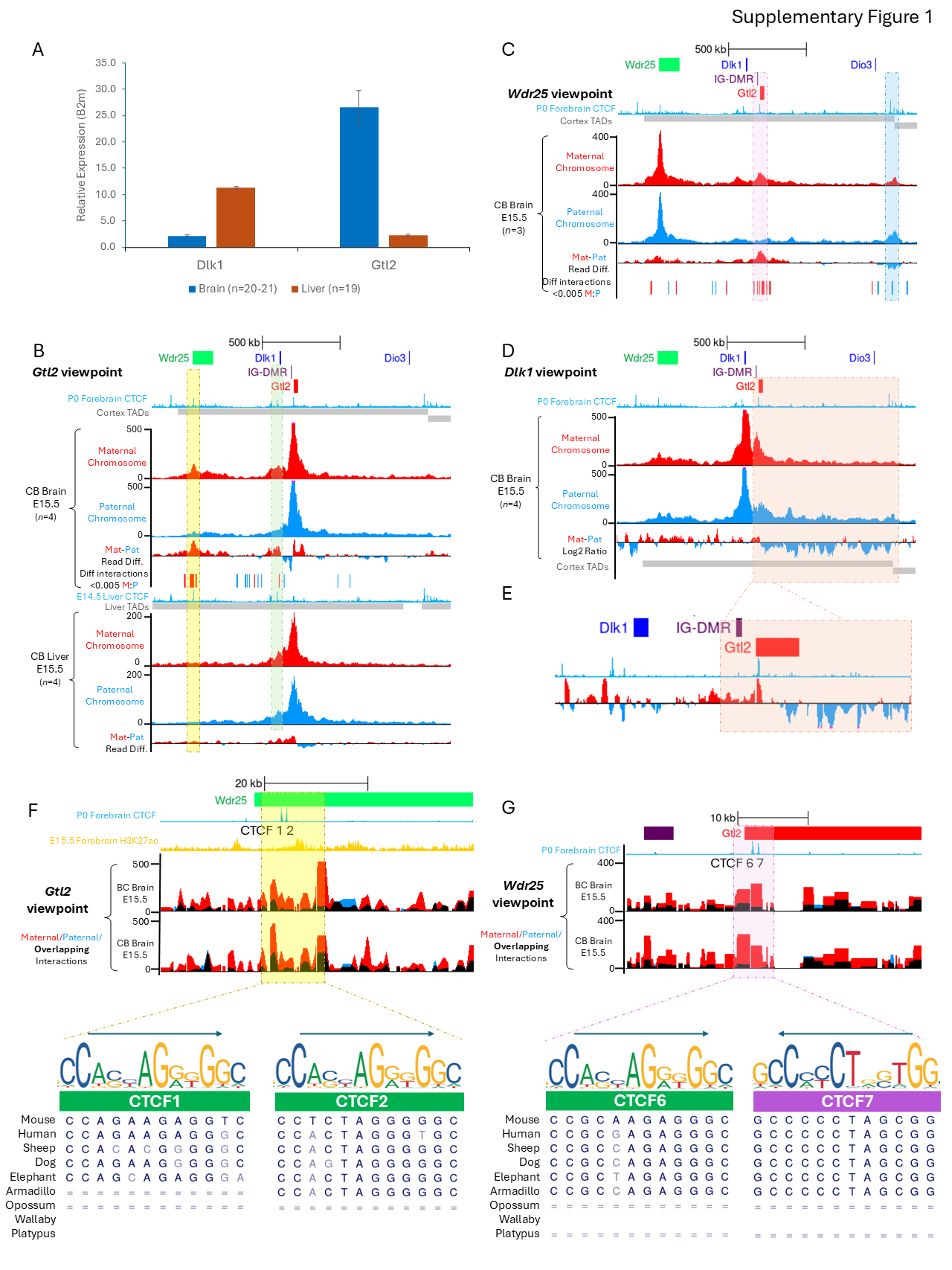

### Supplementary Figure 4.2

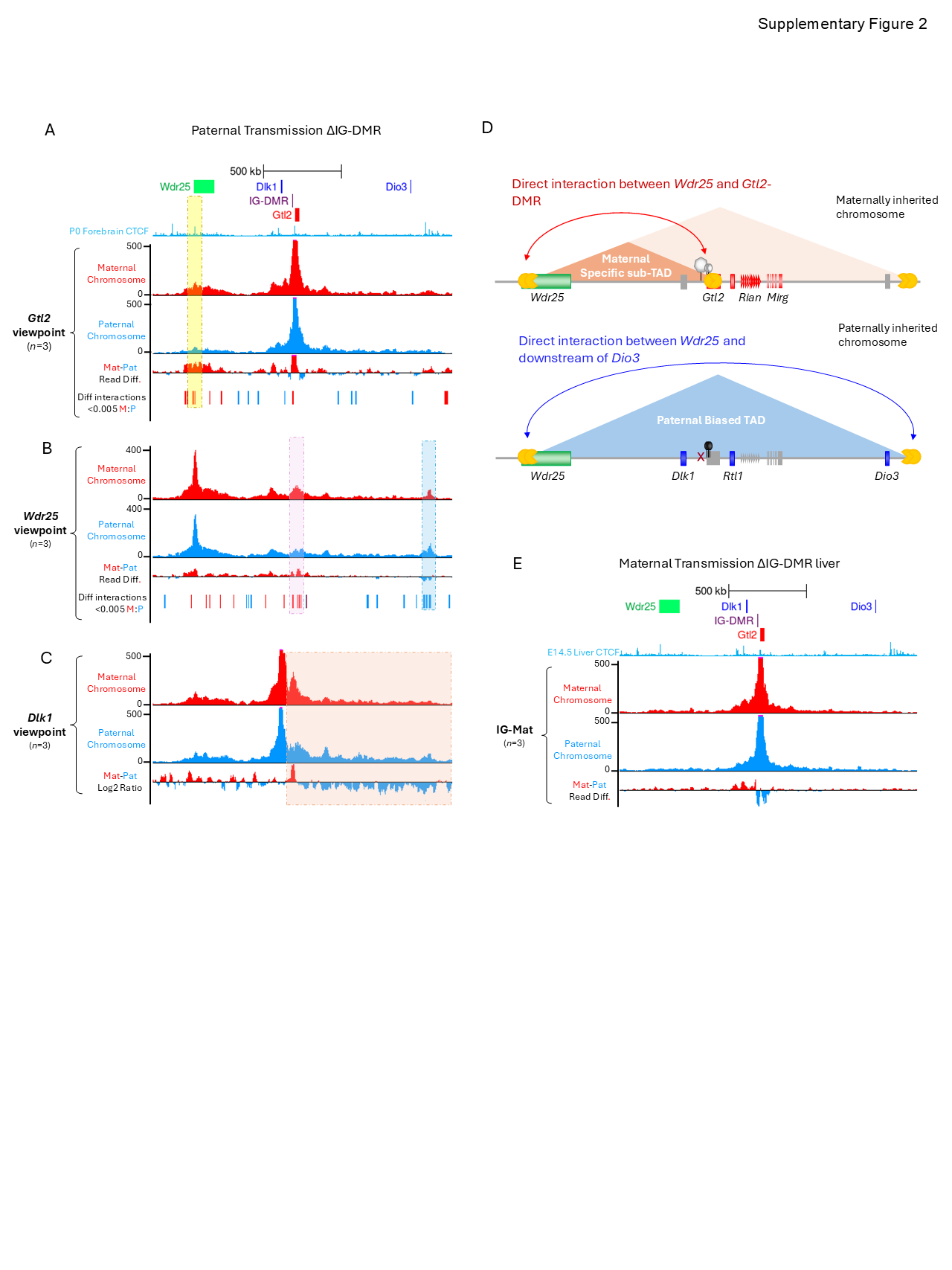

### Supplementary Figure 5

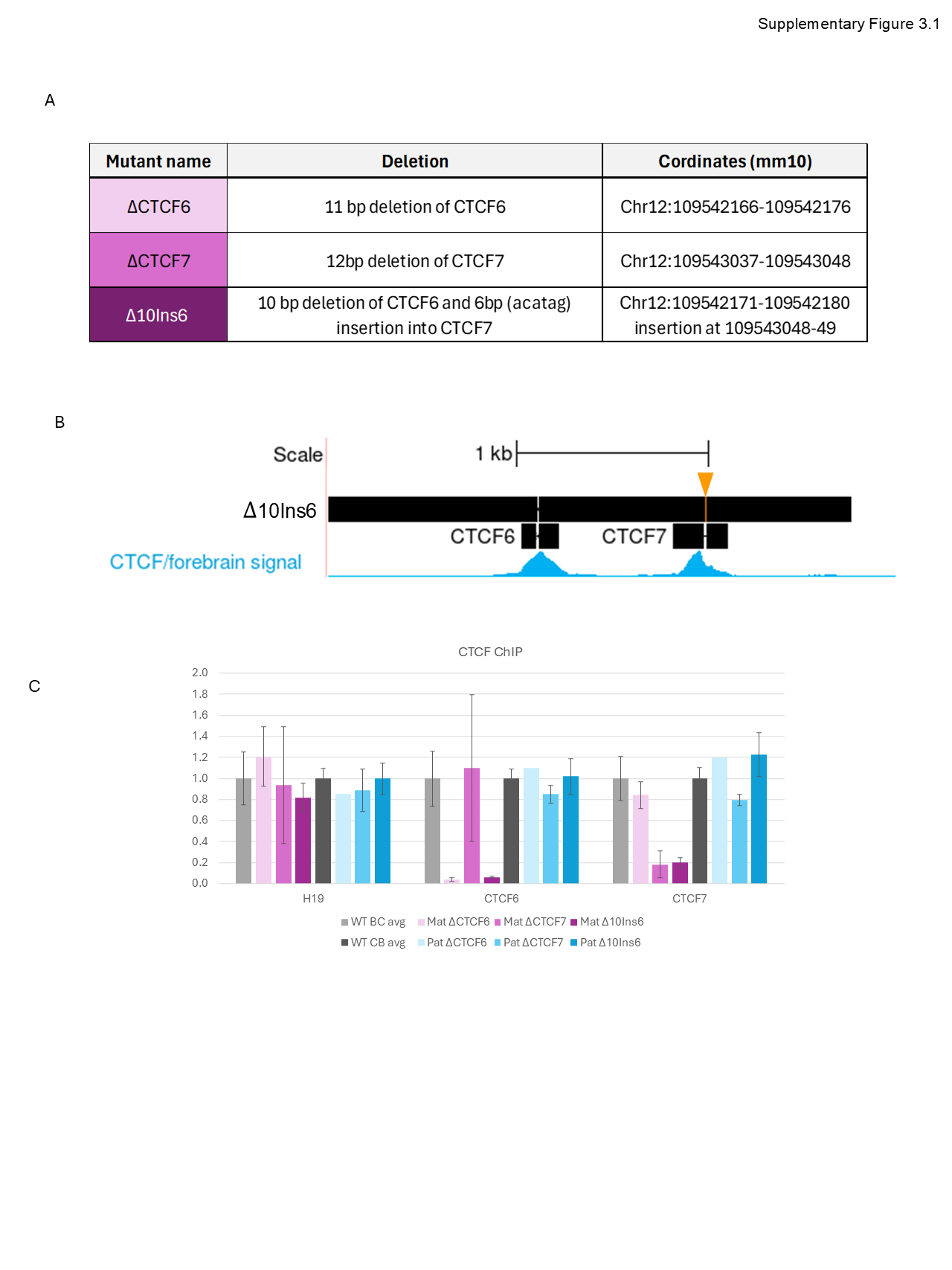
